# Single-nucleus RNA-seq identifies Huntington disease astrocyte states

**DOI:** 10.1101/799973

**Authors:** Osama Al-Dalahmah, Alexander A Sosunov, A Shaik, Kenneth Ofori, Yang Liu, Jean Paul Vonsattel, Istvan Adorjan, Vilas Menon, James E Goldman

**Author notes:** Corresponding Author: James E Goldman, 01-212-305-3554.

## Abstract

Huntington Disease (HD) is an inherited movement disorder caused by expanded CAG repeats in the Huntingtin gene. We have used single nucleus RNASeq (snRNASeq) to uncover cellular phenotypes that change in the disease, investigating single cell gene expression in cingulate cortex of patients with HD and comparing the gene expression to that of patients with no neurological disease. In this study, we focused on astrocytes, although we found significant gene expression differences in neurons, oligodendrocytes, and microglia as well. In particular, the gene expression profiles of astrocytes in HD showed multiple signatures, varying in phenotype from cells that had markedly upregulated metallothionein and heat shock genes, but had not completely lost the expression of genes associated with normal protoplasmic astrocytes, to astrocytes that had substantially upregulated *GFAP* and had lost expression of many normal protoplasmic astrocyte genes as well as metallothionein genes. When compared to astrocytes in control samples, astrocyte signatures in HD also showed downregulated expression of a number of genes, including several associated with protoplasmic astrocyte function and lipid synthesis. Thus, HD astrocytes appeared in variable transcriptional phenotypes, and could be divided into several different “states”, defined by patterns of gene expression. Ultimately, this study begins to fill the knowledge gap of single cell gene expression in HD and provide a more detailed understanding of the variation in changes in gene expression during astrocyte “reactions” to the disease.

## Introduction

Huntington Disease (HD), a neurodegenerative disorder caused by CAG repeats in the Huntingtin gene, leads to the accumulation of the mutant protein and an associated neuronal degeneration and gliosis [34]. Although Huntingtin is expressed in all cell types, the neuropathology of the disease shows substantial variation. For instance, there are caudal-rostral and medial-lateral gradients of severity in the neostriatum [60], and relative sparing of the nucleus accumbens in advanced grade HD [47]. Neuronal degeneration in the isocortex involves the motor and sensory cortices but relatively spares the superior parietal lobule [37], and in some patients, the anterior cingulate cortex is also involved, and cortical layers III, V, and VI are especially vulnerable [48].

Understanding the differences in microenvironment between affected and relatively resistant regions may illuminate the cellular and molecular mechanisms underlying vulnerability and resilience to neurodegeneration. Astrocytes, in particular, are a key to maintaining the neuronal microenvironment, and display regional variation across the dorsal-ventral, rostral-caudal, and medial-lateral axes according to developmentally-determined domains [58]. Additionally, astrocytes exhibit intra-regional heterogeneity. For example, subpial and white matter astrocytes contain more glial fibrillary acidic protein (GFAP) and CD44 than protoplasmic astrocytes but lower levels of protoplasmic astrocyte markers, such as glutamate transporters and glutamine synthetase [54]. Astrocytes in the affected regions of the HD brain show a “reactive” state, generally defined by histochemistry or by an increase in GFAP [51, 61], but also by a decrease in the expression and protein levels of the major astrocytic glutamate transporter, EAAT2 [2,9]. In mouse models of HD, expression of mutant Huntingtin (*mHTT*) in astrocytes leads to decrease in the expression of the glutamate transporter [6], and the failure to buffer extracellular potassium and glutamate leads to neuronal hyper-excitability and a HD-like phenotype, which is reversed by restoration of astrocytic membrane conductance [56]. In addition, astrocytes in HD contribute to an inflammatory environment [16,26], which likely participates in the progression of the pathology. Thus, there is substantial evidence that astrocytes play a primary role in the evolution of HD.

Whereas bulk transcriptome-wide studies have provided important insights into molecular changes in HD [1, 14, 15, 22, 28, 38], bulk samples comprise a mixture of cell types, so astrocyte-specific signatures in HD may be obscured. Single cell RNA sequencing (scRNAseq) is a powerful technique to interrogate cellular heterogeneity [7, 44]. Although brain banks house hundreds of frozen postmortem brain specimens, technical difficulties limit the application of scRNAseq to these samples. However, this is not a limitation of single *<u>nucleus</u>* RNA sequencing (snRNASeq), which accurately elucidates cellular heterogeneity in a manner comparable to whole/cytoplasmic scRNAseq [24], and can be applied to frozen brain tissue [21]. This technique has been used to identify multiple novel, regionally-diversified cortical excitatory and inhibitory neuronal sub-types [23]. Recently, massively parallel snRNASeq using droplet technology revealed cellular heterogeneity in the human postmortem cortex and hippocampus [11, 17]. The disease-specific cell-type specific transcriptional signatures were described in oligodendroglia in multiple sclerosis [18] and multiple cell types in Alzheimer Disease [31] and microglia in Alzheimer disease [39].

In this study, we performed snRNASeq of fresh frozen cingulate cortex from Grade III/IV HD patients and compared the results to those from non-neurological control patients to examine single cell differences in gene expression. We have focused here on astrocytes, although we found significant differences in all cell types. We examined the cingulate cortex, which is often affected in HD patients, because the cortical pathology is less severe than the neostriatal pathology, and because there is little known about cortical astrocyte pathology in HD. Finally, we focused on Grade III/IV to identify astrocyte gene expression changes in an intermediate stage of evolution rather than at an end stage.

Overall, we found substantial heterogeneity in astrocyte signatures, with clear differences in population structure between control and HD tissue samples. In particular, the “reactive” astrocytes in HD could be divided into several different “states”, defined by patterns of gene expression. These ranged from astrocytes that had markedly upregulated metallothionein (MT) and heat shock genes, but had not completely lost the expression of genes associated with protoplasmic astrocytes, to astrocytes that had substantially upregulated *GFAP* and had lost expression of many normal protoplasmic genes as well as the MT genes. The variation in astrocytes is not surprising, given that HD is a degenerative disease that progresses over years and one would not expect all astrocytes to respond synchronously during the course of the disease.

Studies like this one will begin to fill the knowledge gap of single cell gene expression in HD and give us a more detailed understanding of the variation in changes in gene expression during astrocyte “reactions.” Furthermore, network building from these gene sets will help define interacting genes and important regulatory genes behind reactive astrocyte states.

## Methods

### Dissection of the cingulate cortex from frozen tissue

Postmortem anterior cingulate cortex specimens frozen during autopsy from control and grade III/IV HD were obtained from the New York Brain Bank. Four cases (two HD and 2 control) were selected for snRNAseq and 12 cases (6 and 6) for bulk RNAseq, all with RNA integrity numbers of > 7. Cortical wedges measuring ~5 × 4 × 3 mm were dissected on a dry ice cooled stage and processed immediately as described below. A table of the cases and controls used is provided in **Table 1**.

**Table 1.**
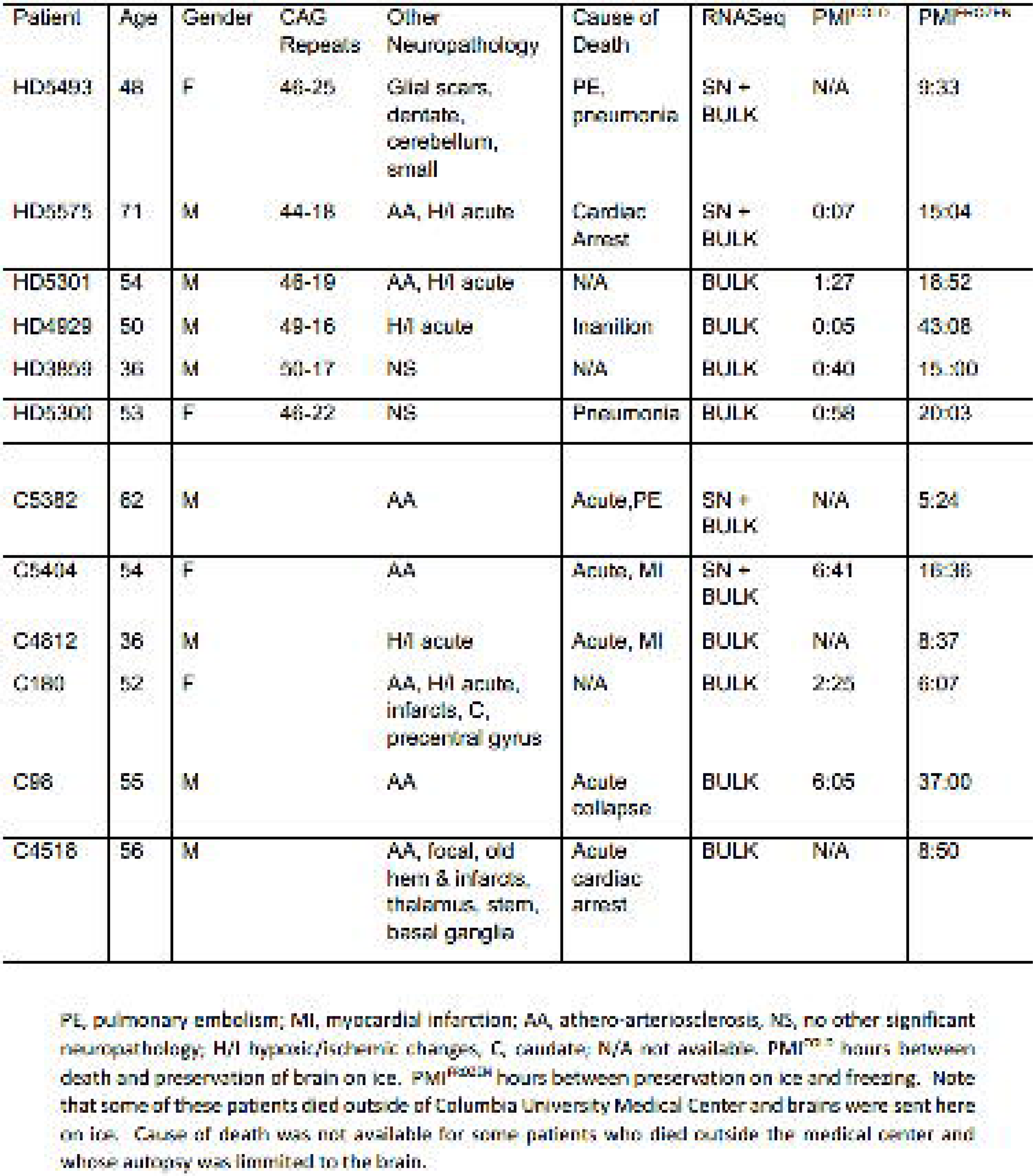
HD and Contron Patient Data

### Single nucleus RNAseq

Nuclei were isolated as described in [21]. Briefly, cortical tissue was homogenized in a Dounce homogenizer with 10-15 strokes of the loose pestle and 10-15 strokes of the tight pestle on ice in a Triton X-100 based, sucrose containing buffer. The suspension from each sample was filtered through a BD Falcon tubes with a cell strainer caps (Becton Dickinson, cat. no. 352235), washed, re-filtered, washed, followed by a cleanup step using iodixanol gradient centrifugation. The nuclear pellet was then re-suspended in 1% BSA in nuclease-free PBS (containing RNAse inhibitors) and titrated to 1000 nuclei/□l. The nuclear suspensions were processed by the Chromium Controller (10x Genomics) using single Cell 3’ Reagent Kit v2 (Chromium Single Cell 3’ Library & Gel Bead Kit v2, catalog number: 120237; Chromium Single Cell A Chip Kit, 48 runs, catalog number: 120236; 10x Genomics).

### Sequencing and raw data analysis

Sequencing of the resultant libraries was done on Illumina NOVAseq 6000 platformV4 150bp paired end reads. Alignment was done using the CellRanger pipeline (10X Genomics) to GRCh38.p12 (refdata-cellranger-GRCh38-1.2.0 file provided by 10x genomics). Count matrices were generated from BAM files using default parameters of the DropEst pipeline [42]. Filtering and QC was done using the scater package [32]. Nuclei with percent exonic reads from all reads in the range of 25-75% were included. Nuclei with percent mitochondrial reads aligning to mitochondria genes of more than 14% were excluded. Genes were filtered by keeping features with >10 counts per row in at least in 31 cells.

### Normalization and data-cleanup

The count matrix was normalized by first running the quickcluster function, then estimating sizefactors by calling scran::computeSumFactors() function with default options and clusters set to clusters identified by calling quickcluster function. Scater::normalize function was then used to generated normalized counts. Of note, batch effects were taken into account when normalizing the data (Scater package in R). Doublet identification was done using scran::doubletCells function with default options, and cells with doublet score of >=1.5 were excluded. A total of 5199 cells passed QC at this point (C5382: 893 nuclei, H5575: 646 nuclei (Batch1), and C 5404: 1096 nuclei, H5493: 2151 nuclei (Batch 2)). We filtered low-quality nuclei using the percentage of the cell-lineage specific genes that are expressed (Identity score – see below). If the cell’s identity score was less than 20%, a nucleus was considered low-quality and was excluded. A total of 4786 nuclei remained after this step.

### Clustering and classification of nuclei

Clustering, cell-lineage assignment, and sub-clustering was done as follows. First, an unsupervised pre-clustering step was performed to split the nuclei into small pre-clusters. Second, the pre-clusters were assigned a class using gene set enrichment analysis and GO term enrichment analysis of the pre-cluster markers. Third, mixed pre-clusters (with mixed enrichment scores for lineage genes) were identified, and the nuclei within these clusters were assigned to new pre-clusters based on their highest identity score (see below). Fourth, the pre-clusters were agglomerated into master classes (Astrocytes, Neurons, Microglia, Endothelial cells, Oligodendrocytes, and Oligodendrocyte Precursor Cells (OPCs). Finally, master classes were sub-clustered using the SC3 clustering algorithm. These steps are described in **Supplementary Figure 7** and in detail below.

### Pre-clustering

First, dimensionality reduction methods including tSNE and PCA (prcomp) were conducted using functions provided in scater package. Briefly, dimensionality reduction by tSNE was done using scater::runtSNE() or runPCA() functions. For runPCA() function, a random seed was set (12345) and theta was set to default (0.5). Multiple perplexity levels were tested –a level of 527 was empirically chosen. We then used an unbiased clustering approach using shared nearest neighbor utilizing the scran::buildSNNGraph() function, with dimensionality reduction set to “TSNE”. Multiple K values were tested (6-16) and the number of pre-clusters generated was examined. K=6 yielded 35 discrete pre-clusters, with the least number of mixed pre-clusters (based on results from the cell classifier-below). K=6 was thus chosen for further clustering. We appreciate that using a highly nonlinear reduction such as tSNE as input for building a SNN graph is not preferable to using PCA. However, our experience is that using tSNE reduced dimensions yielded fewer pre-clusters with mixed signatures of well-studied, known cell classes. Moreover, we used an independent method to ascertain the quality of the pre-clusters (see next section).

We developed a simple algorithm to assign cell classes based on the highest proportion of cell-class specific genes (referred to thereafter as Cell classifier tool). The tool also classifies nuclei into master classes (Neurons, Astrocytes, Oligodendrocytes, OPCs, Microglia, and Endothelial Cells). More specifically, each nucleus is given an identity score that is equal to the percentage of genes that are expressed from a cell-class specific gene list. The gene lists used to classify nuclei are based on the literature [36, 66], modified, and are provided (**Supplementary Table 1**). Each nucleus is given identity scores for all 6 cell types: Neuron, Astrocyte, Oligodendrocyte, OPC, Microglia, and Endothelial. The cell is assigned the class (identity) for which it scores the highest identity score. Additionally, an ambiguity status is assigned based on whether the curve of identity scores is skewed (Unambiguous) towards the highest score versus flattened (Ambiguous). More specifically, if the sum of second and third highest identity scores is higher than 2X the highest identity score, the identity curve is flattened, and the cell is considered ambiguous. This helps identify problematic nuclei in mixed clusters - as explained in the section below.

### Pre-cluster identity assignment

To assign pre-cluster identities, we used multiple approaches to ascertain pre-cluster and nuclei identities. First, we used the mean of normalized counts per gene per cluster to perform gene set enrichment analysis using the gsva R package (’gsva’ method with kcdf=“Gausian” and mx.diff=FALSE). We used cell-type specific genesets based on a literature search (provided in **Supplementary Table 1** – the same as those used for the cell classifier tool). Second, we manually examined the top marker genes that differentiate pre-clusters and determined whether any pre-clusters showed lineage-discordant marker genes. For example, if one pre-cluster was characterized by oligodendrocytic (MBP, TMEM120, MOBP) as well as neuronal (CCK, MAP1B, BEX3, NRGN) or microglial (HLA-B, IBA1, C1QB) genes, a cluster was considered mixed. Finally, we also performed GO term enrichment analysis on the top marker genes and manually examined the terms that distinguished each cluster. Pre-cluster markers were discovered using the scran::findmarkers() function with default options.

The majority of pre-clusters were homogeneous, and showed high enrichment scores in one lineage. However, some pre-clusters, like pre-clusters 2, 7, 27, and 17, were mixed, showing high enrichment scores for microglial and oligodendrocytic genes (2 and 17), or astrocytic and neuronal genes (7 and 27). Thus, we used supervised classification, as above, and re-assignment of the nuclei in the ambiguous clusters based on the maximum score assigned by the cell classifier scores (cluster_max_score). This resulted in the following pre-clusters: Microglia_r2_17, Oligodendrocyte_r2_17, Astrocyte_r2_17, Neuron_r17_2, Endothelial_r2, Astrocyte_r7_1, Astrocyte_r27_2, Neuron_r7_1, and Neuron_r27_2. In addition, cluster 31 showed mixed neuronal and microglial enrichment scores, but based on the cell classifier scores the majority of cells 51/59 were designated as neurons, and only one cell was called microglial. (Astrocyte-6, endothelial cells-0, Microglial-1, Neuron-51, Oligodendrocyte -1). Thus, this cluster was renamed as 31_Neuron. Note, for endothelial cells, only supervised assignment was able to detect a discrete cluster.

Afterwards, clusters of the same lineage were conglomerated into master classes: Astrocytes (1064 nuclei), Endothelial cells (19), Microglia (147), Neuron (2085 nuclei), Oligodendrocytes (1230 nuclei), and OPC’s (241 nuclei).

For neuronal clusters, a minority of nuclei expressed astrocytic genes despite the neuronal identity being the highest as assigned based on the cell-classifier results. These cells had ambiguous scores per the cell classifier. Thus, all neuronal nuclei that were classified as ambiguous were excluded (219 nuclei), leaving 1866 nuclei for downstream analysis (to do SC3). *GFAP* and *AQP4* genes were excluded from the analysis of neuronal nuclei (the counts were re-normalized without reads from *GFAP/AQP4* genes).

### Consensus clustering using SC3 for sub-clustering

To determine objectively the optimal way to cluster the astrocytic, oligodendrocytic, and neuronal nuclei, we used consensus clustering as described in the SC3 package [19]. Values for K (number of clusters in K-mean clustering) were empirically tested and silhouette widths average values for the resultant clusters were examined. For astrocytes K=6 was chosen as silhouette values were all above 0.5 (except cluster 2, an HD astrocytic cluster with both *GFAP* expression and MT gene expression. We decided to keep this cluster as it made biological sense. Cluster markers (generated through the SC3 pipeline) were then generated using pairwise t-test. To find marker genes (SC3 package), a binary classifier was constructed as informed by the mean cluster expression values for each gene. Prediction accuracy was tested using the area under the receiver operating characteristic (AUROC) curve. Wilcoxon signed rank test p-values were assigned per gene. Here we used (AUROC) > 0.65 and with the p-value < 0.05. Cluster markers were also generated using pair-wise t-test (as per scran library in R). Note, for the astrocytic sub-cluster gene list used to classify astrocytic nuclei, the following options were using for the scran::findmarkers() function: log fold change 0.5 and markers were detected in “any” direction. A selected list of astrocytic sub-cluster top gene markers and cell-type cluster markers are provided in **Table 2** and **Supplementary Table 1**, respectively. These lists were generated as follows: For each cluster, the log-fold change values for each gene were summed across the other clusters. The genes were then ranked in descending order by these sums. Genes with one or more negative log-fold changes or FDR-corrected p values≥0.05 were excluded from the marker list, even if the sum of the log-fold changes was high. This imparts specificity to the markers selected for each cluster.

**Table 2.**
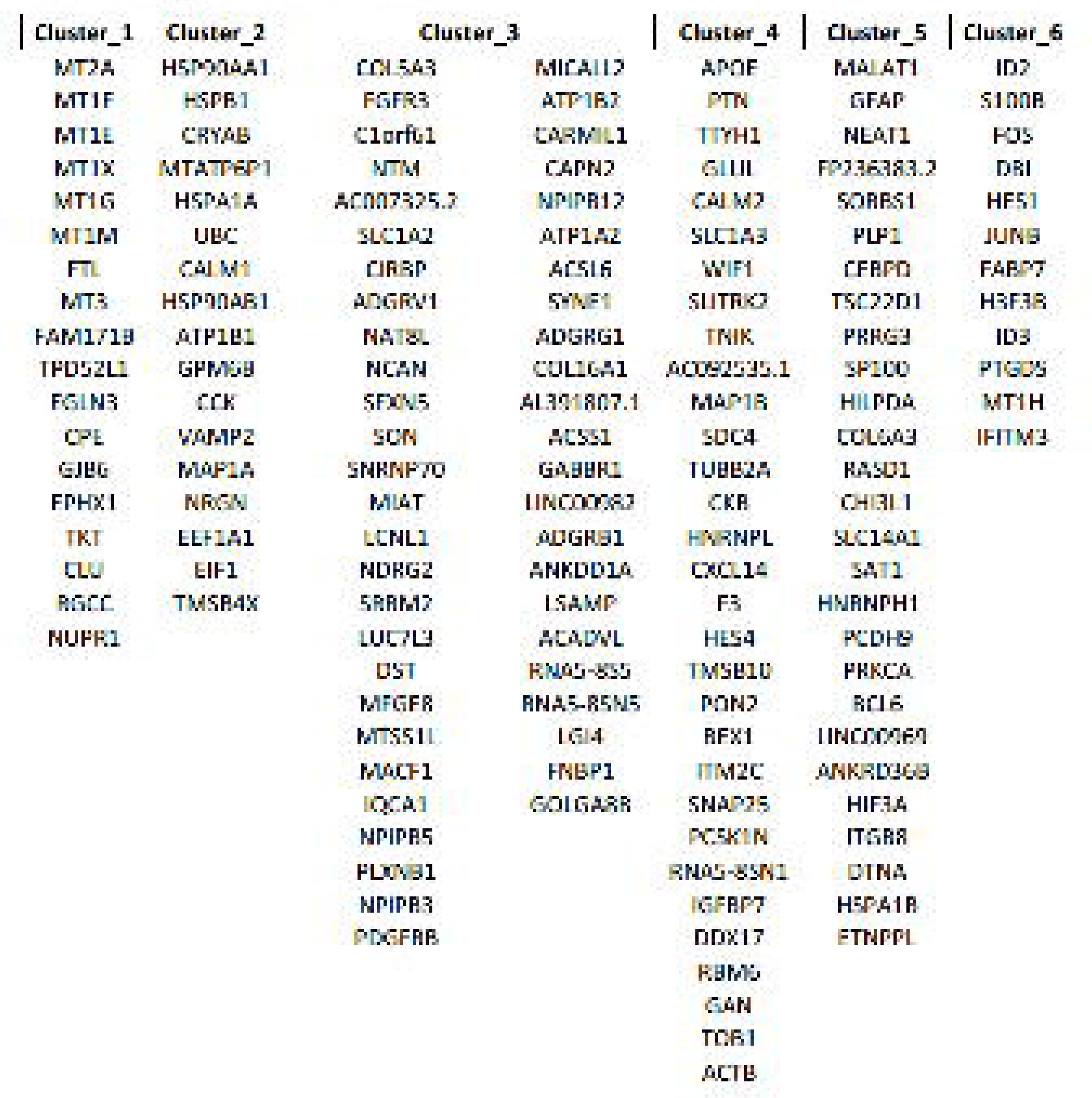
Astrocytic sub-cluster markers

### Supervised classification of astrocytes

To classify astrocytic nuclei into astrocytic states, we used monocle 2.0 [57]. We used the following rules to assign astrocytes into four states based on log-transformed expression values: Quiescent-SLC1A2>= 2, MT2A <4, and GFAP < 3; State-1Q: SLC1A2>= 2, MT2A >=4, and GFAP < 3; State-2R: SLC1A2< 2, MT2A >=4, and GFAP >=3; and State-3R: SLC1A2< 2, MT2A <4, and GFAP >=3. Events that did not meet any of the conditions, or met more than one condition were classified as unknown or ambiguous, respectively.

### Differential gene correlation analysis and gene Multiscale Embedded Gene Co-Expression Network Analysis

Differential gene correlation analysis and gene network analysis were done in DGCA [35] and MEGENA [52] R packages, respectively. In brief, the count matrix was filtered to keep the top 5% most highly expressed astrocytic genes by average using the filtergenes() function with the following options: filterTypes=“mean” and filterCentralPercentile = 0.95. The resultant matrix had a total of 856 genes. The DGCA pipeline was performed using the ddcorAll() function with the following options: adjust = “perm”, nPerm = 100, nPairs = 1000. Visualization was done in ggplot2 [64]. Identification of astrocytic gene modules was done in MEGENA R package with default options (min size =10). GO term enrichment analysis of genes in the modules was done using the moduleGO() function in DGCA R package with a p value set at 0.05. Gene set variation analysis of module genes was done using the GSVA() R package as described below.

### Gene set variation analysis (GSVA)

The average normalized count per gene per cluster was calculated. The resultant cluster-wise count matrix was used as input to the GSVA pipeline [12]. Gene sets used for various tests are provided in the supplementary material (**Supplementary Table 1**). The options used for performing the GSVA pipeline are as follows: method= ‘gsva’, kcdf=“Gaussian”, mx.diff=FALSE. Heat maps were generated using the heatmap.2 in R function from the package gplots [63] and scores z-scaled were indicated.

### Bulk RNAseq

RNA was extracted from cingulate cortical specimens using Trizol™ method. RIN values were determined, all samples had values>7.0. Sequencing was done on Illumina Hiseq 2000™ platform. Raw reads were aligned to the reference (GRCh38.p12) using STAR aligned [8]. Ribosomal RNA reads were removed using SortMeRNA [20]. For data analysis was done in R v 3.5 in Linux or Windows 10 environment. Genes with fewer than 10 reads per row were excluded. Principal component analysis of normalized variance stabilized counts (product of DESeq2::varianceStabilizingTransformation) in FactoMineR R package [25] revealed that the first two components explained 47.4% of the variance in the data, and that the variance was captured entirely in the first 11 components. Condition (HD versus control) was the variable that is best correlated with PC1 (eta2=0.62, cos2=0.88). After variance stabilization, library size was not correlated with PC1 (correlation coefficient 0.08). For differential gene expression and statistical analysis, EdgeR [33, 46] with incorporating age and gender into the design matrix. The likelihood ratio test (LRT) method was used with an adjusted p value of 0.05 and an absolute log fold change threshold set at 1.4. Principal component analysis and visualization were done using DESeq2 package [30]. Sample distances were calculated from normalized counts (DESeq2) using the Manhattan metric using the dist() function from base package in R (v3.5.2). pheatmap from pheatmap package was used for visualizing utilizing the ward method for clustering [45]. Gene Ontology term analysis was done in gProfiler web platform (https://biit.cs.ut.ee/gprofiler/gost) by supplying an ordered query and determine statistical significance using the Benjamini-Hochberg FDR method (p<0.05). The terms shown in the Figures are selected based on ordering the results based on negative_log10_of_adjusted_p_value followed by the ratio of the shared of number of genes enriched in a term to that of the total number of genes in the GO term (desc(intersection_size/term_size)).

### Quantitative PCR

Total RNA was extracted from brain specimens using TRIzol reagent (Invitrogen, Carlsbad, CA, USA). RNA concentration and purity were determined using NanoDrop (Thermo Scientific™, MA). A bioanalyzer was used to determine RNA integrity. RNA was converted to cDNA using High capacity RNA-to-cDNATM kit (Thermo Fisher Scientific, Applied BiosystemsTM,MA). Specific qPCR primers were designed using Primer3 and from Primer Bank. Each 15 μL qPCR reaction contained 5ng of cDNA, 7.5 μL of 2- SsoAdvanced™ Universal SYBR® Green Supermix (Biorad, PA), 400nM of each primer and nuclease free water. The qPCR plates were read on a Mastercycler® RealPlex2 (Eppendorf, NY). The reactions were done in triplicates. Relative gene expression was calculated using the delta delta Ct Pfaffl method with UBE2D2 and RPL13 as reference genes (Pfaffl MW. 2001). Statistical analysis was done using one-tailed Mann Whitney U test. The *GFAP* primers were AGGTCCATGTGGAGCTTGAC (forward) and GCCATTGCCTCATACTGCGT (reverse) and *ACTNB* primers were CTGGAACGGTGAAGGTGACA (forward) and AAGGGACTTCCTGTAACAATGCA (reverse) [65].

### Human brain tissue processing, histology, *in situ* hybridizations, and immunohistochemistry

Standard H&E and Cresyl violet histochemical stains were done in the histology core at the Department of Pathology and Cell Biology at Columbia University. The cases and controls used are provided in **Table 1**. Standard chromogenic and fluorescent Immunohistochemistry was done as described previously in Sosunov A et al. 2014 in paraffin-embedded formalin-fixed tissue sections or fresh frozen sections briefly fixed in 4% PFA, for 10 min (4° C) in 4% PFA in PBS. Paraffin sections after deparaffinization were treated with Antigen Unmasking Solution according to the manufacturer’s recommendations (Vector Laboratories, Burlingame, CA). GFAP and Huntingtin dual immunohistochemical (IHC) stains were done using standard DAB and alkaline phosphatase dual staining on a Leica Bond™ auto-stainer. The following antibodies and dilutions were used GFAP (rabbit polyclonal, 1:1000, DAKO Z0034) or chicken polyclonal, Abcam Cat# ab4674, 1:300), CD44 (rat monoclonal, 1:100, Millipore A020), glutamine synthetase (GS) (mouse monoclonal, 1:1000, Transduction Laboratories 610518), C3 (rabbit monoclonal, 1:200, Abcam Ab20999), metallothionein (MT) (mouse monoclonal, 1:200, Abcam ab12228), HTT (mouse monoclonal, 1:2000 Millipore MAB5492 1:2000), ALDH1L1 (mouse monoclonal, Encor Cat# MCA-2E7, 1:100), IBA1 (Wako # 019-19741 Rabbit polyclonal, 1:300-1:500), LN3 (mouse monoclonal, 1;75, MP Biomedicals LLC, #69303). For fluorescent IHC, secondary antibody conjugated to fluorophores: anti-mouse Alexa Fluor 488 and 594, anti-rabbit Alexa Fluor 488 and 594, and anti-chicken Alexa Fluor 620; all from goat or donkey (1:300, ThermoFisher Scientific, Eugene, OR) were applied for 1 hr RT. *In situ* hybridization was done using RNAscope™ multiplex fluorescent v2 (ACDbio cat no 323100) per the manufacturer’s protocol in 5-micron paraffin-embedded, formalin-fixed tissue sections. We used custom designed probes for GFAP and PLP-1 (cat no.’s 584791-C3 and 564571-C2, for GFAP and PLP-1, respectively). The probes were designed to cover all transcripts. We used an available probe for MBP (Cat. no. 573051). Images were taken on a Zeiss 810 Axio confocal microscope. Brightfield and chromogenic images were taken on an Aperio LSM™ slide scanner at 20X (Brightfield).

## Results

### Major gene expression alterations in the HD cingulate cortex

To explore gene expression alterations in the cingulate cortex, we analyzed bulk RNA expression profiles from six grade III and grade IV HD and six non-neurologic controls (**Table-1**). Analysis of sample distance showed HD samples clustered together, except for one case of Juvenile onset HD (H3859) (**Figure 1A**). A recent report suggests that in juvenile HD the cerebral cortex is largely unchanged compared with controls (Tereshchenko A. et al 2019). We next performed differential gene expression analysis from bulk RNAseq specimens, and identified 3165 downregulated genes and 1835 upregulated genes at a Benjamini-Hochberg adjusted false discovery rate of 0.05 (**Figure 1B**). Reactome pathways and GO terms enriched in the upregulated genes revealed an enrichment in HD of multiple immune response genes including complement, toll-like receptor signaling, and Interleukin pathways, featuring IL-13 and IL-10 (**Figure 1D**). In addition, genes involved in responses to metal ions, metal sequestration, and metallothionein binding were also enriched in HD. GO term analysis of genes reduced in HD revealed that the majority of these genes were associated with neuronal identity or function (**Supplementary Figure 5D**). This is further discussed in the Supplementary Results. A full list of the top differentially upregulated and downregulated genes in HD is provided in **Supplementary Table 5**.

**Figure_1.**
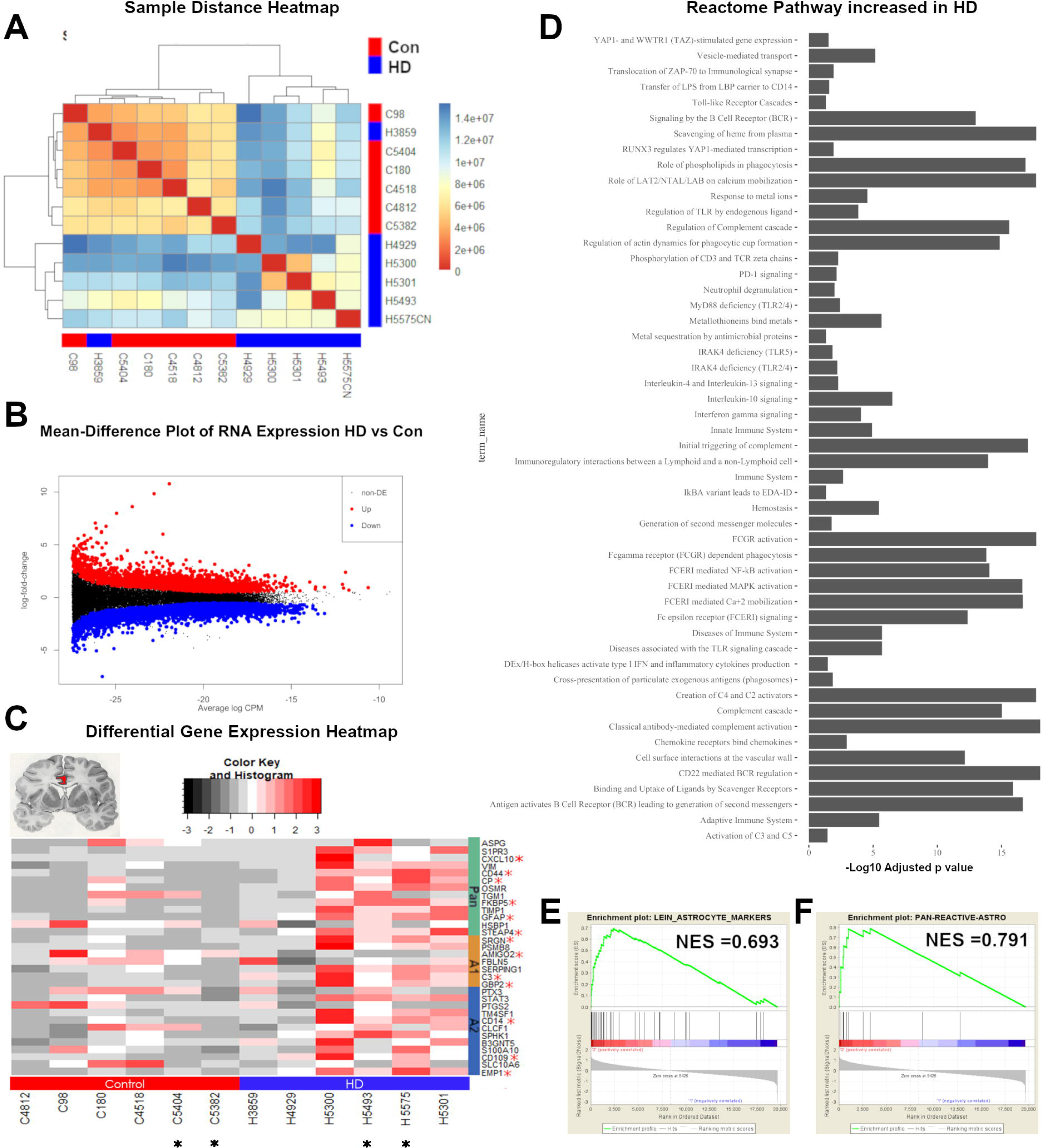
Bulk_RNASEQ Transcriptomic analysis of Huntington disease cingulate cortex. Total RNA sequencing was done on 6 grade III/IV and 6 control cingulate cortices. **A)** Sample distance (Manhattan method) heatmap clustered using the Ward method. **B**) Mean-Expression plot showing log2 fold-change (LFC-HD versus control) on the y-axis, and mean normalized counts on the x-axis. Significantly differentially expressed genes are shown in red and blue for upregulated and downregulated genes, respectively. **C**) Differential gene expression heatmap of select astrocytic genes as described in Liddelow et al. [26], with controls (Con) denoted by the red bar, and HD by the blue. Asterisks next to gene names indicate significance (Benjamini-Hochberg adjusted p value < 0.05 and absolute log fold change >= 1.5). A1, A2, and pan-reactive astrocytic genes are denoted. Asterisks below case numbers indicate which cases were selected for single cell nuclear RNAseq. **D**) Representative GO ontology term analysis showing significantly increased Reactome pathways in HD cases from bulk RNAseq (using the gProfiler web-platform, Benjamini-Hochberg adjusted p-values set at < 0.05). **E-F**) Gene set enrichment analysis of select astrocytic genes (Astrocyte markers Lien ES et al. 2007 and Astrocyte_differentiation -GO0048708). Normalized enrichment scores (NES) are shown.

Next, we examined astrocytic genes that were differentially expressed in HD. We chose to look at the expression of astrocyte genes from Zamanian et al. and Liddelow et al. [26, 66], described as A1 (neurotoxic), A2 (neuroprotective), and pan-reactive astrocytic genes (**Figure 1C**). We found that a subset of A1, A2, and pan-astrocytic genes were significantly increased in HD, including *GFAP*, *CD44*, *OSMR*, *FKBP5*, *STEAP4*, and *CXCL10* for pan-reactive genes, *C3*, *SRGN*, and *GBP2* for A1 genes, and *EMP1*, *CD14*, and *CD109* for A2 genes. In contrast, the gene *AMIGO2*, an A1-specific gene, was reduced in HD. We next performed geneset enrichment analysis of select astrocytic genesets (astrocyte differentiation and astrocyte markers, [27] and found them enriched in HD (**Figure 1E-F**). Of note, there was significant heterogeneity in differential gene expression of astrocytic genes among HD cases. For example, RNA expression profiles from the aforementioned juvenile HD case (H3859) as well as another grade IV HD case (H4929) showed little upregulation of astrocytic genes (**Figure 1C**). We investigated this further using immunohistochemistry for astrocytic markers GFAP, GS, and ALDH1-L1 (**Supplementary Figure 1G**). The results showed no qualitative changes in pattern of staining or densities of astrocytes in the cingulate cortex in case H3859 compared to non-neurologic controls (**Supplementary Figure 1G**). Taken together, the data show major gene expression changes in the HD cingulate cortex, involving upregulation of immune genes as well as astrocytic genes.

### Single nucleus RNAseq from control and HD cingulate cortices identifies major cell types

The results of the bulk RNAseq showed heterogeneity within the HD cases, with some cases not showing significant differences in expression of astrocytic genes compared to controls (T4929 and T3859). Moreover, a subset of A1 “neurotoxic”, A2 “neuroprotective”, and pan-reactive genes were increased HD, and it was not clear whether this represents a coexistent increase in A1 and A2 astrocytes, versus increased expression of A1 and A2 genes in the same cells. To address the issue of astrocytic heterogeneity, we selected 2 HD (grade III) and 2 control cases, extracted nuclei from the cingulate gyrus, and then performed snRNAseq using the 10X Genomics Chromium™ droplet-based single cell RNAseq platform (**Figure 2A**). 4786 nuclei passed quality control measures, of which 1989 were control and 2797 were HD nuclei, as shown in the t-distributed stochastic neighbor embedding (tSNE) plot in **Figure 2B**. We used both unsupervised clustering and supervised classification to identify cell types, including neurons, astrocytes, oligodendrocytes, OPCs, microglia, and endothelial cells (**Figure 2C**). Scaled normalized gene expression in individual nuclei per cell-type displayed in a heatmap show distinct gene expression profiles among nuclei in different cell types (**Figure 2D**). Gene-set variation analysis of the average normalized gene expression per cell type against cell-type specific gene-sets derived from thorough assessment of the literature and genes from Gill B. et al. [10] (**Supplementary Table 1A**) show high enrichment of cell-type specific gene-sets in the classified cell types (**Supplementary Figure 2A-B).** The proportions of each cell type per condition are shown in **Supplementary Figure 2C**. While the proportions of astrocytes (24% controls and 21% HD) and microglia (3% each) were comparable, the proportions of neurons (53% controls and 37% HD) and oligodendrocytes (15% controls and 33% HD) were divergent. We cannot ascertain whether this is due to technical versus biological effects. The numbers and proportions of nuclei per cell type per case are shown in **Supplementary Figure 2D-E**, respectively.

**Figure 2:**
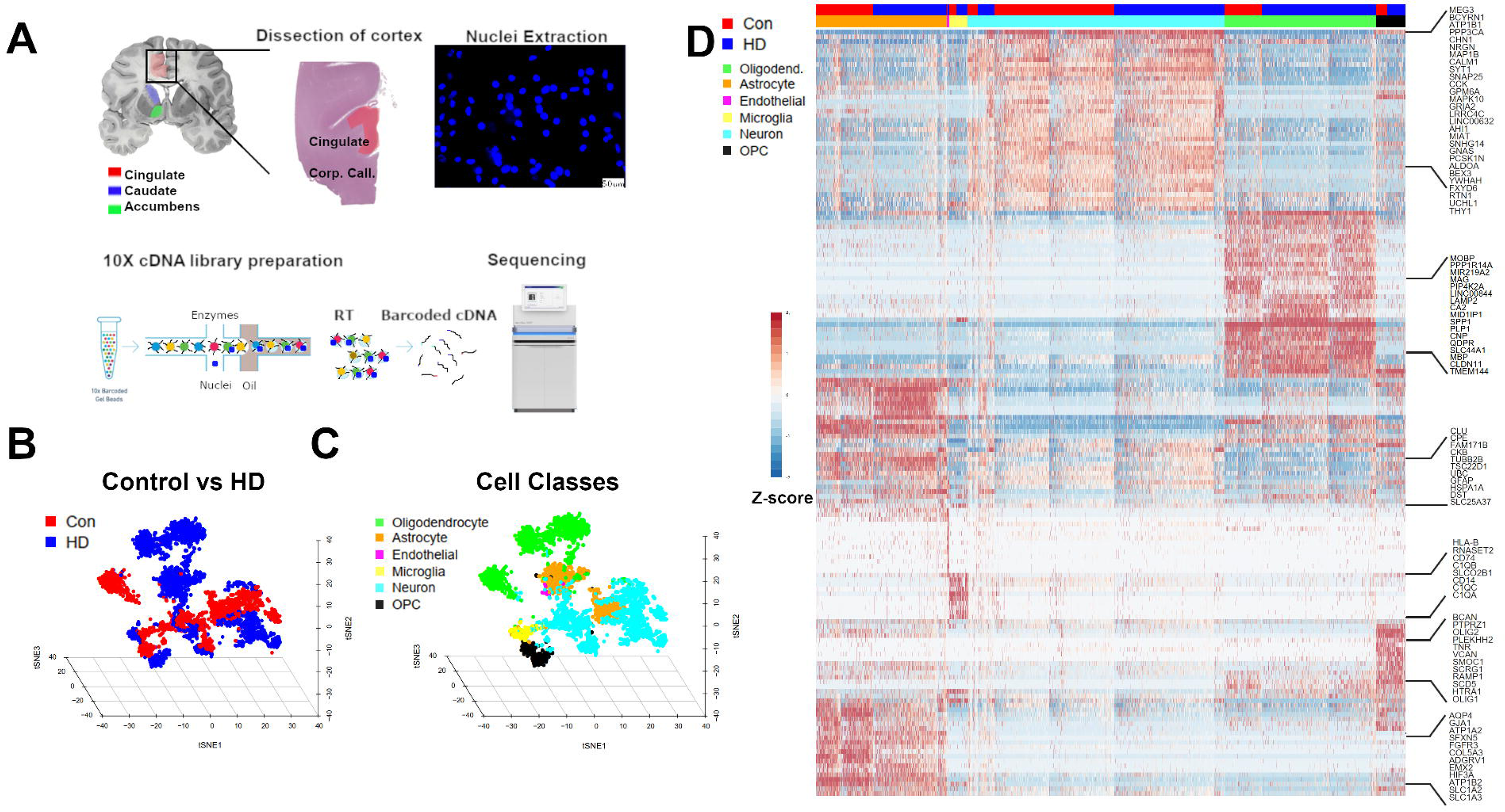
Single cell nucleus RNAseq of the cingulate cortex in Control and HD. **A**) Experimental scheme; first, cingulate cortex was dissected, nuclei were extracted and visualized using DAPI nuclear stain under a fluorescence microscope to ascertain membrane integrity. The nuclei were subjected to 10X chromium single cell RNAseq workflow involving encapsulation of nuclei in oil droplets along with enzymes and barcoded beads, followed by cDNA synthesis and library preparation, and finally, sequencing. **B**) Three-dimensional t-distributed stochastic neighbor embedding (tSNE) plot showing individual nuclei (Points, n = 4786) colored as red (Control) or blue (HD), reduced into three dimensions. **C**) Three dimensional tSNE plot showing the classification of the nuclei into neurons (Cyan), Oligodendrocytes (Green), Astrocytes (Orange), Oligodendrocyte precursor cells (OPC-Black), Microglia (Magenta), and endothelial cells (Purple). **D**) Gene expression heat map of z-scaled normalized counts showing nuclei (Columns) and specific cell-type markers (Rows), a subset of which are shown on the right. Condition (Con versus HD) and Cell-types are color-coded on the top as in panels B-C.

### Astrocytes in the HD cingulate cortex

To investigate astrocytic changes in HD in the cingulate cortex, we focused on astrocytic nuclei (1064) and projected these nuclei into the tSNE space (**Figure 3A**). Control and HD astrocytic nuclei were separated in the tSNE space. We next performed sub-clustering using consensus k-means clustering in the SC3 package in R and identified six clusters, three control and three HD astrocytic, with minimal overlap between the conditions (**Figure 3B** and **Supplementary video 1**). We next identified cluster-specific markers using the SC3 package pairwise gene Wilcoxon signed test with p.value of 0.05, Holm’s method for p.value adjustment, and area under receiver operator curve (AUROC) 0.65 (**Figure 3C**). We then examined differentially expressed genes in HD astrocytes versus control astrocytes. 1645 genes were downregulated in HD while 607 genes were increased (exact p value and Adjusted Benjamini-Hochberg adjusted p.value < 0.05). Analysis of enriched gene ontology terms and Reactome pathways showed the top gene ontology molecular function terms included terms related to heat shock protein binding, unfolded protein binding, MHC binding, and metal ion binding, while enriched Reactome pathways included response to metal ions, metallothionein binds metals, and Heat-shock factor 1 (HSF-1) dependent transactivation (**Figure 3D**). In contrast molecular function gene ontology terms enriched in downregulated astrocytic genes included terms related to symporter activity, amino acid binding, and neurotransmitter:Sodium symporter, while enriched Reactome pathways included Notch signaling and cholesterol biosynthesis (**Figure 3E**). A list of differentially expressed genes in HD versus control astrocytes as well as the full results of GO term enrichment analysis are provided in **Supplementary Table 2A-B**. A selected list of astrocytic sub-cluster marker genes is provided in **Table 2**. The full results of differential gene expression between each sub-cluster against all other sub-clusters is provided in the Supplementary data.

**Figure 3:**
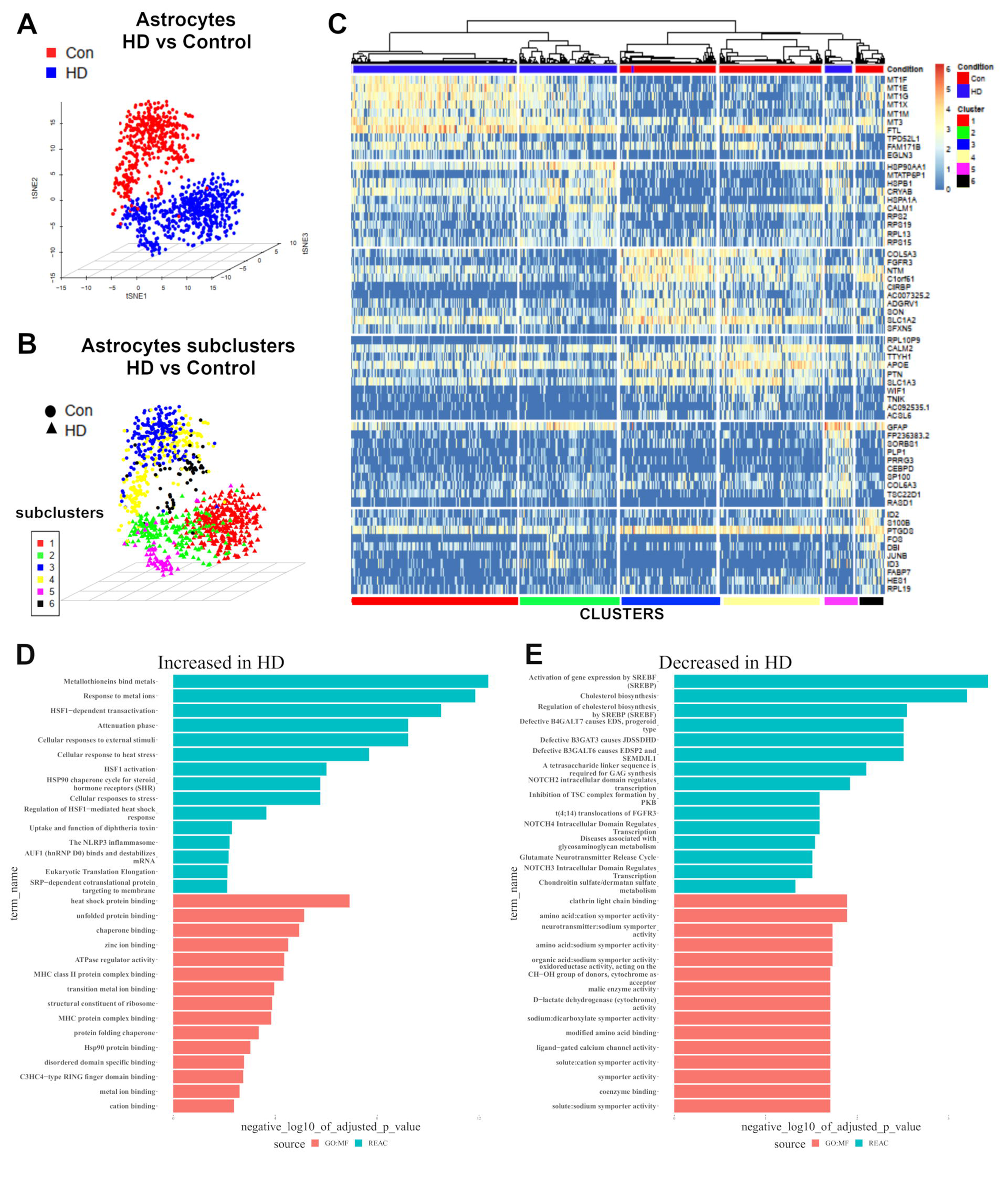
Transcriptomic analysis of astrocytic nuclei in control and HD. **A**) Three-dimensional t-distributed stochastic neighbor embedding (tSNE) plot showing astrocytic nuclei (n = 1064 −469 control and 595 HD) colored as red (Control) or blue (HD), reduced into three dimensions. **B**) Three dimensional tSNE plot showing the classification of the astrocytic nuclei into 6 sub-clusters using consensus k-means clustering (sc3 package). **C**) Gene expression heat map of cluster markers showing nuclei (Columns) and specific cell-type markers (Rows). Condition (Con versus HD) and Cell-types are color-coded on the top and bottom, respectively. Cluster-specific gene markers were identified using Wilcoxon signed rank test comparing gene ranks in the cluster with the highest mean expression against all others. p-values were adjusted using the “holm” method. The genes with the area under the ROC curve (AUROC) >0.65 and p-value<0.01 are shown as marker genes. **D-E**) GO term analysis of differentially expressed genes in HD versus control astrocytes identified using EdgeR likelihood ratio test with an adjusted p.value of 0.05. Significantly enriched GO terms at Benjamini-Hochberg false discovery rate of 0.05 were identified. Selected Reactome pathways and Molecular function GO terms which are increased in HD astrocytes (**D**) and decreased in HD astrocytes (**E**) are shown.

Since Liddelow et al. [26] showed that astrocytes in the HD caudate nucleus expressed markers of a putative neurotoxic state (A1 state), we compared the cingulate cortex to the caudate using an antibody to C3 – a marker of the A1 astrocytic state. We confirmed that numerous cells morphologically consistent with astrocytes in the caudate nucleus of HD grades (III and IV) were C3 positive compared to controls. Double immunostaining for C3 and GFAP (**Supplementary Figure 3D**) as well as C3 and LN3 (microglial marker), showed immunopositivity for C3 in GFAP-positive astrocytes, and minimal immunopositivity in microglia (**Supplementary Figure 3C, D**). In contrast, immunostaining in the cingulate cortex showed C3 labeling of neurons in controls and HD cases, but no astrocyte labeling, with no clear differences (**Supplementary Figure 3A-B**). Together, these findings confirm previous reports of the A1-astrocytic marker C3 immunopositivity in striatal astrocytes in HD, but, in contrast, show minimal astrocytic labeling with C3 in the HD cingulate cortex.

### Multiple genes are differentially correlated in HD and control cortical astrocytes

We examined the gene-expression heatmap for the 6 astrocyte clusters (**Figure 3C**). Clusters 1, 2, and 5 were from HD cortex, while 3, 4, and 6 were from control cortex. It is apparent that the control clusters 3 and 4 express high levels of genes associated with protoplasmic astrocytes, like *FGFR3*, *GLUL*, and *SLC1A2*, although there are differences from cluster to cluster and from cell to cell, and control cluster 6 shows high levels of *GFAP*. The HD clusters 1 and 2 express high levels of MT genes (*MT1F*, *MT1E*, and *MT1G*.), as well as *GFAP*, although MT genes were generally more highly expressed in cluster 1 than in cluster 2 or cluster 5 and *GFAP* was more highly expressed in cluster 2 and 6 than in cluster 1. We provide the full differential gene expression between astrocytic clusters in a **Supplementary Data file (Supplementary_Data_2)**.

Given this heterogeneity of gene expression, we set out to identify how astrocytic genes are co-regulated. We performed differential gene correlation analysis between the top 5% most highly expressed astrocytic genes on average using the DGCA R package. The results revealed the MT genes were highly correlated with each other, and the correlation was significantly stronger in HD astrocytes than in control astrocytes (**Figure 5A**). Moreover, there were differences in the correlation of many gene pairs between control and HD astrocytes. For instance, multiple MT genes were negatively correlated with *GFAP*, including *MT1G*, *MT1F*, MT1E, and *MT2A* in HD astrocytes. However, these gene pairs were either not correlated or only minimally positively correlated with *GFAP* in control nuclei (**Figure 5A** and **Supplementary Table 3**). As an example, a scatter plot showing the expression of the *GFAP* and *MT1G* in control and HD astrocytes reveals that the two genes are negatively correlated (**Figure 5B**); with Pearson correlation coefficient of 0.21 in HD and 0.18 in control astrocytes (adjusted p value of difference <0.0001). In contrast, MT genes *MT1F* and *MT2A* were positively correlated with genes associated with protoplasmic astrocytes including *SLC1A3* in HD, but negatively correlated in control astrocytes (Pearson correlation coefficients of 0.14 (HD) and −0.14 (Control) for *MT2A* - adjusted p value of difference < 0.01, and 0.1 (HD) and −0.14 (Control) for *MT1F* -adjusted p value of difference < 0.05 – **Supplementary Table 3**).

These analyses of HD and control astrocytes reveal a heterogeneity of gene expression phenotypes in both the HD and control populations that would not have been detected without snRNA-seq. The heterogeneity also adds a layer of complexity both to the populations of astrocytes in control cortex as well as to the populations of astrocytes reacting to a disease.

### Gene co-expression network analysis in HD and control astrocytes

Next, we aimed to identify gene networks in which subsets of astrocyte genes were linked to each other. We used the Multiscale Embedded Gene Co-Expression Network Analysis (MEGENA, [52]). A usual application involves building gene networks in the control condition, and examining how the network structure changes in the test condition. However, given that control and HD astrocytes clustered separately, and there were significant gene expression changes between the two conditions, we decided to use all astrocytes from both conditions for gene network analysis. This allowed discovering both condition-specific and shared gene networks. Application of the algorithm produced 15 gene clusters (modules). The network hierarchy structure is shown in **Figure 5C**. The modules on the same arm of the hierarchy tree shared many genes. For example, modules 3 and 10, modules 22 and 26, as well as modules 9 and 20, shared many genes (**Supplementary Table 4A-B**, which details all the module genes and hub genes). Examples of modules are shown in **Figure 5D**.

We then correlated each module with one or more of the 6 astrocyte clusters. To indicate the functional characteristics of the modules, we used Gene Ontology term enrichment analysis to ascribe “Molecular Function” gene ontologies that were enriched in each of the modules. Modules 9 and 20 were mostly enriched in the HD cluster 1 (compare **Figures 5E** and **F**) and have genes associated with molecular function gene ontology terms like amyloid beta binding and low-density lipoprotein particle receptor binding – genes known to be associated with reactive astrocytes in Alzheimer’s disease [31, 50]. Module 7 was also most enriched in cluster 1 and has genes enriched for GO terms related to metal binding (see above regarding MT genes). Modules 6 and 16 are enriched in the HD cluster 5, and include genes enriched for GO terms related to epithelial-mesenchymal transition, RNA-mediated gene silencing, and DNA binding (**Figure 5E-F and Supplementary Table 4C-D)**. Modules 4, 12, 22, and 26 are enriched in control cluster 3 and have genes enriched in GO terms related to regulation of neurotransmitter levels, autophagy, sodium ion transport (Module 12), and cell-cell signaling. Module 5 was most enriched in control cluster 6 and has genes involved in protein transport, localization, and biosynthesis. Module 17 was enriched in control cluster 4 and has genes enriched for GO terms related to nucleotide triphosphatase, pyrophosphatase, and hydrolase activity. Shared between HD cluster 2 and control cluster 4 are Modules 3 and 10, which have genes enriched for GO terms related to aerobic respiration, nucleotide metabolism, and ATP biosynthesis. Of interest, *Module 13* was enriched in control astrocytic clusters Astrocyte_3 and Astrocyte_4 (**Figure 5F**), and showed enrichment for GO terms relating to fatty acid biosynthetic process, fatty acid binding, long-chain fatty acid biosynthetic process, and prostaglandin biosynthetic process (**Supplementary Table 4C-D**). Conversely, *Module 20* was enriched in HD cluster Astrocyte_1 and less so in control cluster Astrocyte_4. This module was enriched for GO terms relating to lipid metabolic process, phospholipid binding, and lipid transport (**Supplementary Table 4C-D**). The top gene ontologies per module are provided in **Supplementary Table 4D**. Thus, in most cases, the network modules appear distinct for either control or HD astrocytes, although there is some overlap. Molecular function analysis further highlights the heterogeneity of both control and HD astrocyte populations.

### Validation of astrocytic gene expression changes in HD

The snRNASeq analysis revealed significant differences in astrocyte gene expression between HD and controls. We wanted to validate these gene expression differences and therefore conducted immunohistochemical and *in situ* hybridization studies in HD and control cingulate cortices (**Figure 4**). To quantify astrocytic activation in the HD cingulate cortex we measured the cell density of GFAP immunopositive cells (**Figure 4A**). We found that the number of GFAP immunopositive cells was increased in grade III/VI HD compared to control (**Figure 4C** and see **Supplementary Methods**). We also confirmed that the *GFAP* transcript levels were indeed increased in the HD cingulate cortex using rt-qPCR (**Figure 4D**). Moreover, we quantified the optical density of the GFAP signal in fluorescently immune-stained sections as a surrogate for protein content in each astrocyte (**Supplementary Methods**). We found that GFAP levels as measured by GFAP signal optical density was increased in HD astrocytes (34.98 +/− 1.92 arbitrary units; mean +/− standard error of the mean (sem)) compared to controls (19.83 +/− 1.95 arbitrary units; mean +/− sem, p < 0.001).

**Figure 4:**
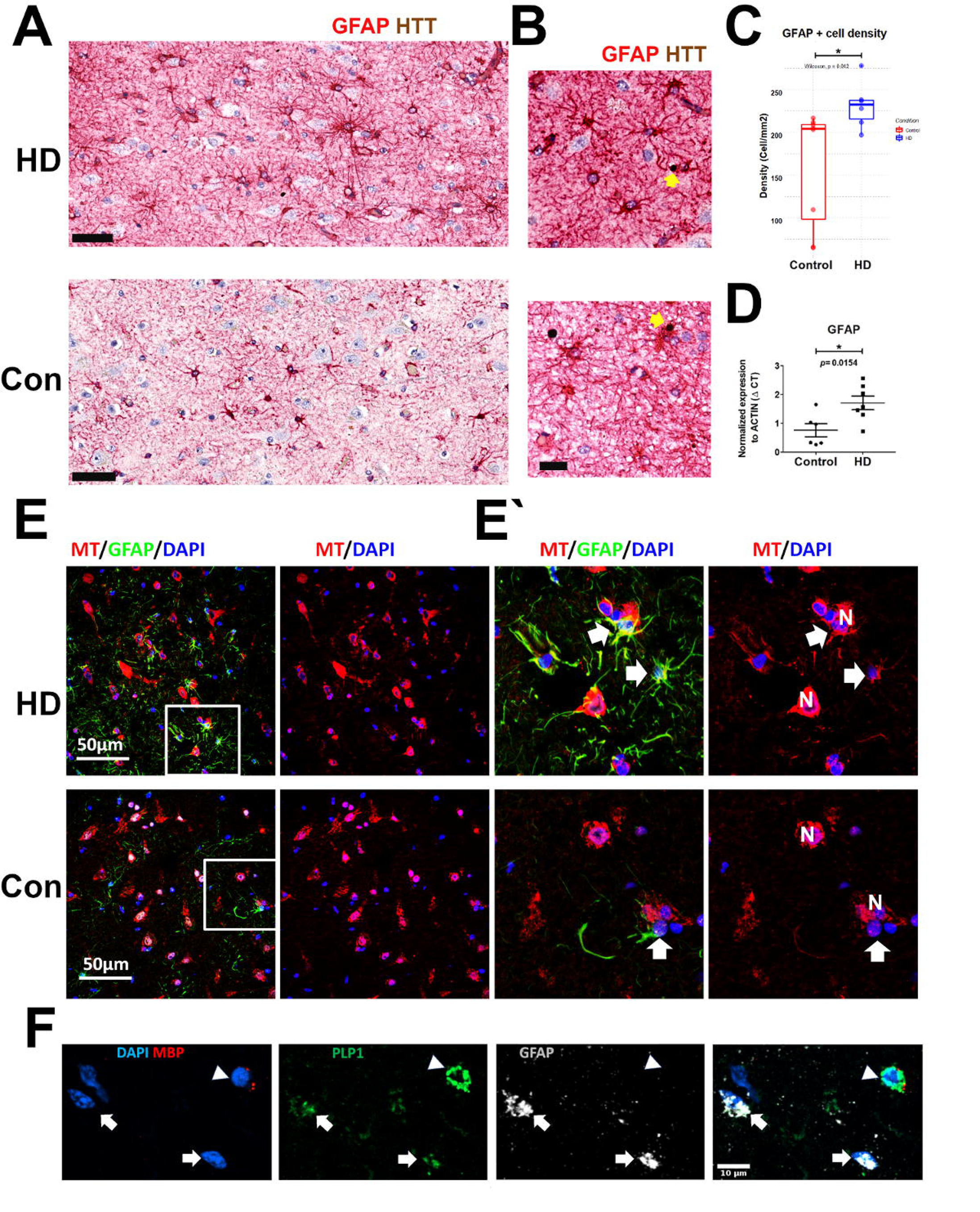
Validation of astrocytic activation in HD. **A**) Representative micrograph of GFAP-HTT dual immunohistochemical stain showing increased GFAP immunoreactivity in the HD cingulate cortex compared to control (scale bar = 50um, GFAP in red & HTT in brown). **B**) Representative images showing accumulation in HTT in the neuropil and in astrocytic in two HD cases (yellow arrows - scale bar = 20um). **C**) Quantification of A, showing increased GFAP immunoreactive cell density in the HD cingulate cortex, n=8 for control and 6 for HD (grade III and IV), P value = 0.012 (one-sided t-Mann-Whitney U test). **D**)Real-time quantitative PCR showing relative expression of GFAP transcript in grade III/IV HD (n=7 for HD and 6 for control) Delta CT value normalized to B-Actin transcript levels are shown. P value = 0.0154 (one-sided t-test). **E**) Representative fluorescent immunohistochemical stains showing GFAP (red), metallothionein (MT-green), and DAPI stained nuclei (blue) from a representative control and grade IV HD case. There is increased GFAP immune-positive cells in layers V/VI of the HD cingulate cortex, and large proportion of these astrocytes are MT positive, compared to control (scale bar = 60um).60um). **F**) In situ hybridization (RNAscope™) showing probes for MBP (red), GFAP (white), PLP1 (green), and DAPI-stained nuclei (blue). Note the co-localization of GFAP and PLP-1 (white arrows). An MBP positive PLP-1 positive Oligodendrocyte is indicated by the arrow head. (scale bar = 10um).

Because HD is caused by HTT mutations, we examined HTT transcript levels in astrocytes in control and HD patients. We found that levels of HTT RNA were reduced in HD astrocytes (**Supplementary results**). We next wanted to ascertain whether HTT accumulates in HD astrocytes. Therefore, we double stained sections with the GFAP antibody and an antibody for wild-type HTT to detect HTT aggregates in astrocytes. We found that HTT does indeed aggregate in HD astrocytes (**Figure 4B**) mainly in layers V and VI. These data suggest that astrocytic reactivity in HD may at least be partly cell-autonomous, or possibly that astrocytes phagocytose HTT aggregates from the surrounding neuropil.

Next, we validated the increase in MTs in HD astrocytes, using immunohistochemistry in the HD cingulate cortex versus controls (**Figure 4E**). The results showed more HD astrocytes co-expressed MT; with 45.57 +/− 25.58% of GFAP positive or ALDH1L1 positive astrocytes co-labeling with MT compared to 10.24 +/− 7.64% of control astrocytes (mean +/− standard deviation, p = 0.043, one-sided t-test). Additionally, we found that levels of MT were also increased in HD astrocytes compared to control astrocytes. More specifically, we measured MT fluorescent signal (optical density) and found it was increased in HD astrocytes (24.33 +/− 1.76 arbitrary units; mean +/− sem) compared to controls (12.3 +/− 1.14 arbitrary units; mean +/− sem, p < 0.001). These results show that both more HD astrocytes expressed MT, and that they expressed higher levels of MT, confirming the snRNAseq results. Note that these MT+/GFAP+ astrocytes are likely to represent HD cluster 2, which, as noted above, expresses the highest levels of MT genes.

To identify cluster 1 astrocytes, which express little GFAP but relatively elevated levels of MT, we used a pan-astrocytic marker, ALDH1L1, in triple immunohistochemical preparations. We showed that astrocytes in HD cluster 1 are indeed identifiable in the HD cingulate cortex, with low to undetectable GFAP levels and immunopositivity for MT (**Supplementary Figure 8**). These are contrasted to cluster 2 astrocytes which show dual-immunopositivity for GFAP and MT (**Supplementary Figure 8 and Figure 4E**).

Interestingly, we noted co-expression of proteolipid protein (*PLP-1*), a myelin gene, and *GFAP* in HD cluster 5 (**Figure 3C**). We verified this in the tissue sections using *in situ* hybridization. The results showed that while *PLP-1* and *MBP* were co-expressed in oligodendrocytes in the cortex and the white matter, *GFAP* and *PLP-1* were co-expressed in a subset of astrocytes in the HD cortex (n=5 cases examined, **Figure 4F**). In contrast, only one case from 4 controls showed rare *PLP-1* co-expression with *GFAP* in the cortex. Notably, gene expression analyses did not reveal other oligodendrocyte genes, including *OPALIN, MOG, MAG, CA2, CD82, OMG MYRF, SOX10*, and *NAFSC*, implying that the GFAP+ astrocytes were not upregulating a coordinated transcriptional program of oligodendrocyte maturation.

### Three astrocytic states of reactivity in HD

A simplified view of astrocytic gene expression profiles reveals patterns based on the expression of protoplasmic astrocytic genes, reactive astrocytic genes, and MTs. We performed a supervised classification of astrocytes based on expression of *MT2A, GFAP*, and *SLC1A2*. The results show three distinct astrocytic states in HD and control astrocytes. Astrocytes with low levels of *MT2A*, low *GFAP* and high *SLC1A2* were classified as Quiescent. Astrocytes with high levels of *MT2A*, low *GFAP* and high *SLC1A2* were classified as state-1Q – for quiescent astrocytes with early reactive features. Astrocytes with high levels of *MT2A*, high *GFAP* and low*SLC1A2* were classified as state2-R – for reactive astrocytes with high *MT*s. Astrocytes with lower levels of *MT2A*, high *GFAP* and low*SLC1A2* were classified as state3-R – for reactive astrocytes with lower *MT*s (**Figure 6A**). As expected, unclassified states are also seen. These represent astrocytes that did not meet any of the classification criteria (“unknown” type of astrocytes), or astrocytes that met more than one category (“ambiguous”). Possibly, these may represent transitional states. We then examined the relative preponderance of these astrocytic states in control versus HD (**Figure 6B**). We found that the Quiescent state is abundant in control astrocytes (49.3%), but low in HD (1.6%). In contrast, states1-Qand 2-R were more abundant in HD (37.4% and 14.0% in HD versus 21% and 1.4% in control). State3-R was relatively low in both conditions, but more abundant in HD than control (7.9% versus 2.5%).

The states of astrocyte reactivity correspond to specific astrocytic clusters (**Figure 6C**). For example, in contrast to the Quiescent astrocytic state, which was comprised of almost entirely control clusters, astrocytic state-1Q was mixed, with a major contribution from HD cluster 1 and smaller contributions from other astrocytic clusters. Moreover, state-2-R was primarily composed of HD clusters 1 and 2, while state-3-R was largely comprised of HD clusters 2 and 5. In summary, we find three reactive astrocytic states based on the expression of *GFAP*, *SLC1A2*, and *MT*s (**Figure 6C**). These states relate to reduction of expression of protoplasmic genes (e.g. *SLC1A2, FGFR3, GLUL*), and increased MTs in one reactive subset of astrocytes versus a preponderance of *GFAP* expression in another reactive subset.

It is difficult to relate these states to the A1 versus A2 astrocytes. For starters, none of the “states” or clusters in the cingulate cortex showed *C3* expression. Moreover, only 25 of 39 astrocytic genes described by the Barres group were detected by scnRNAseq, and only 3 were significantly increased in HD (GFAP, SRGN, and CD14). Taken together, we cannot ascribe an A1 versus A2 phenotype to cingulate HD astrocytes.

### Gene expression analysis of non-astrocyte cells

Although we have focused on astrocytes in this report, snRNAseq provided gene expression patterns of all other cell types, neurons, microglia, oligodendrocytes, oligodendrocyte precursors, and endothelial cells. We found differences in gene expression patterns for both neurons and microglia that distinguished HD from control cortex, and present these findings in the Supplement. For oligodendrocytes we have evidence that the heterogeneity of cells of the oligodendrocyte lineage is increased in HD, with a shift to less mature phenotype (manuscript in preparation).

## Discussion

### The HD cingulate cortex contains multiple reactive states of astrocytes

One of the most notable findings in this study is that there are multiple states of astrocyte reactivity, rather than a single reactive state. This is not surprising, given that HD is a long-term, progressive disease in which the neuropathology does not evolve in synchrony. HD cluster 1 showed high levels of *MT* gene expression, and relatively low *GFAP* expression. In contrast, HD cluster 5 showed high *GFAP* expression and relatively low *MT* gene expression. Intermediate levels of *MT*s and *GFAP* were the hallmark of HD cluster 2. In all HD astrocytes, the mRNA levels of glutamate transporters *SLC1A2* and *SLC1A3* expressed in quiescent astrocytes were reduced, implying that HD astrocytes downregulate genes central to some critical astrocytic functions. It is tempting to speculate that these states reflect a temporal sequence of astrocyte reactivity, and that one progresses to another over time, so that the highest *GFAP* astrocytes that have lost many normal genes as well as *MT*s represent an end stage of reactivity. We do not have definitive evidence that this is so. However, when we analyzed striata from these same brains we saw a greater proportion of astrocytes in this possible end stage state (unpublished data), given the far high degree of neuronal degeneration in striatum than cingulate. Additional evidence from examining HD cases with different grades of severity as well as evidence from experimental models such as astrocyte lineage tracing experiments in conditional HD mice models are needed to query this possibility further. Alternatively, difference in astrocytic reactive states may reflect spatial differences in astrocyte localization in cortical layers with variable susceptibility to neurodegeneration, such as susceptible layers III, V, and VI versus resilient layer II [48].

The presence of high expression of *MT* genes, as well as anti-oxidant genes in HD cluster 1 may suggest that this astrocyte “state - State 1Q” is protective. In fact, these astrocytes may be protective of neurons and also of themselves, an “astro-protective” state. Resolving how this state relates to the A2 state is an interesting matter, however, it is not possible to evaluate this putative relationship without additional experimentation. Are both states underpinned by the same molecular signaling pathways, such as STAT3 activation for example? Or are these states divergent? Such questions are intriguing and will be the subject of future work.

### The control cingulate cortex also contains a heterogeneity of astrocytes

Not only did we find a heterogeneity of astrocyte phenotypes in the HD cortex, we also found variation in astrocyte populations in the control cortex, based on the gene expression profiles. For example, astrocyte cluster 6 appears different from both 3 and 4 in that it has higher levels of *GFAP, CRYAB, FOS, JUNB, ID2* and *ID3* and lower levels of *GLUL, SLC1A2*, and *SLC1A3*, indicating that astrocytes of this group have downregulated protoplasmic astrocyte genes and upregulated “reactive” genes. Clusters 3 and 4 also show differential gene expressions. The heterogeneity of astrocyte gene expression must be validated and explored in depth in the future. However, it does raise the possibility that there is a significant amount of astrocyte heterogeneity in the control cortex. We do not know if this heterogeneity is fixed, or whether gene expression variations are transient, depending on such factors as neuronal activity or local blood flow, for example. Furthermore, the presence of astrocytes with “reactive” markers in a “control” cortex may well reflect an individual’s age, past medical history or terminal illness, even in patients without histories of neurological diseases.

### HD astrocytes upregulate metallothionein genes

*MT* genes are thought to confer neuroprotection. Levels of MTs are increased in multiple injuries including ischemia, heavy metal exposure, infection, and neurodegeneration in AD [49]. Mathys and colleagues investigated the frontal cortex of human late onset Alzheimer disease using snRNAseq and showed expression of *MT2A*, *MT1G*, and *MT1E* in astrocytic cluster Ast1 [31]. Mice deficient in *MT1/2* show impaired repair and wound healing after freeze injury, with increased microglia/macrophage infiltration, astrocytosis, and apoptosis [41]. Neuronal survival was compromised in *MT1/2* knockout mice after middle cerebral artery occlusion [62]). Conversely, mice overexpressing *MT1* showed increased resilience to ischemic damage in the same model of injury [59]. The evidence suggests that upregulation of *MT*s maybe a protective response to injury and we propose that it may be so in HD astrocytes. As we have noted above, upregulation of *MT*s may not only be neuroprotective but also astro-protective. A recent study of HD is also consistent with upregulation of *MT* genes. Lin, L et al. [29]. examined transcriptional signatures in the motor cortex (BA4) of grade 2-3 HD showed that GO terms involved in Response to Cadmium Ion (GO0071276), Zinc ion (GO0071294, GO0010034) were increased, while Cholesterol synthesis was reduced (e.g. GO0006695). These data are consistent with our results in HD astrocytes and suggest that upregulation of *MT*s in reactive astrocytes is a pan-disease reactive response. It is known that oxidative stress is a characteristic of the HD brain [53]. The fact that MTs are involved in combating oxidative stress, and that they are upregulated in HD astrocytes, impart a potential neuroprotective role for this phenomenon, perhaps an A2-like state.

On another note, our data highlights fatty acid synthesis and lipid metabolism as key processes that are differentially regulated in HD astrocytes. We find that HD astrocytic clusters had downregulated genes that are associated with fatty acid and prostaglandin biosynthesis. Indeed, the gene PTGSD (Prostaglandin D2 Synthase) and FASN (Fatty Acid Synthase) were reduced in HD astrocytic clusters (**Figure 3C and Supplementary Table 2C**). Moreover, cholesterol biosynthesis was reduced in HD astrocytes (**Figure 3E**). In contrast, lipid binding and metabolism were processes enriched in genes characteristic of HD astrocytic cluster 1 (Module 20 - **Figure 5F and Supplementary Table 4C-D**).

**Figure 5:**
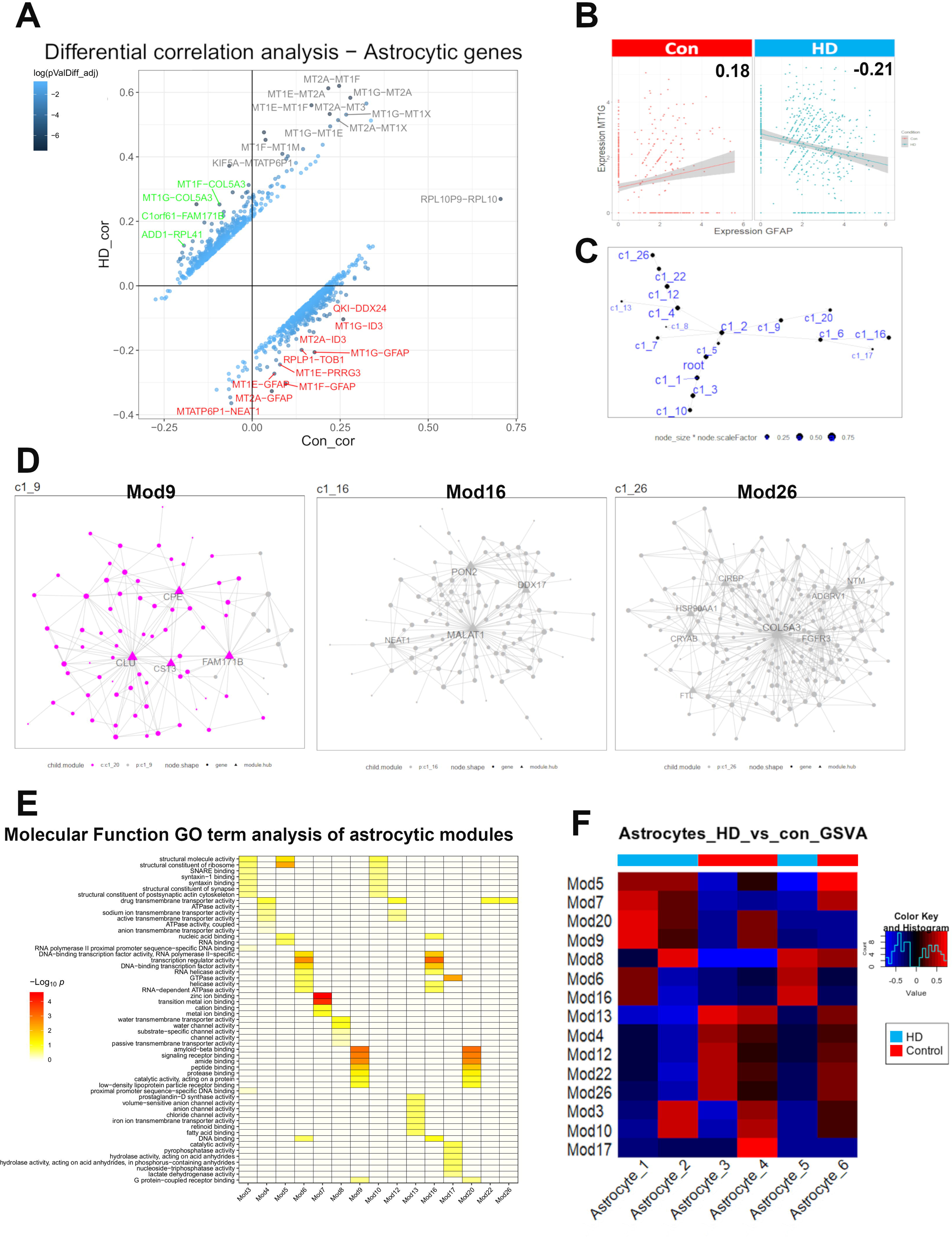
A deeper look into astrocytic gene regulation in HD. **A)** Differential gene correlation analysis of the top 5% of genes in astrocytic nuclei by mean normalized expression (856 genes). Pearson correlation coefficients are shown and empirical p-values were calculated by a permutation test (100 permutations) in DGCA package in R. Top gene-pairs that are significantly differentially correlated between HD and control astrocytes are labeled. **B)** Scatterplot of normalized expression of GFAP and MT1G (A representative metallothionein gene) in Control (left panel) and HD (Right panel) astrocytic nuclei. Pearson correlation coefficients are shown and are significantly different between control and HD. **C)** Hierarchy structure representing the astrocytic gene modules/networks (labeled here as c1_3, c1_5, … etc) as the output using Multiscale Embedded Gene Co-Expression Network Analysis (MEGENA). Clusters on the same line are more related to each other than clusters on different lines. **D)** Representative gene modules/networks, illustrating modules 9, 16, and 26. Hub genes are shown as triangles, genes as nodes, different colors represent different networks in the same graph (Left panel – Mod9). **E)** Gene Ontology term enrichment analysis showing representative “Molecular Function” gene ontologies enriched in astrocytic gene modules. **F)** Gene set variation analysis showing enrichment scores of modules in different astrocytic clusters, condition is shown as colored bars on the top.

### Astrocyte Modules

Analyzing astrocytic gene expression in terms of gene modules provides a useful way to examine correlated genes and hub genes. The latter may be prioritized in functional studies such as overexpression and knockdown/knockout studies. We constructed the modules using all astrocytes, from control and HD brains. Had we constructed modules from HD and control astrocytes separately, the network structures would have been different. For example, module_26, a module harboring many protoplasmic genes, would likely have been devoid of *CRYAB*, a gene upregulated in reactive states. The strength of our approach; combining both control and HD astrocytes, is that it allows us to discover both HD and control modules as well as shared modules, such as module_3 (shared between clusters Astrocyte_2 and Astrocyte_4). Genes in this module relate to oxidative phosphorylation and Wnt signaling (**Figure 6C**).

**Figure 6:**
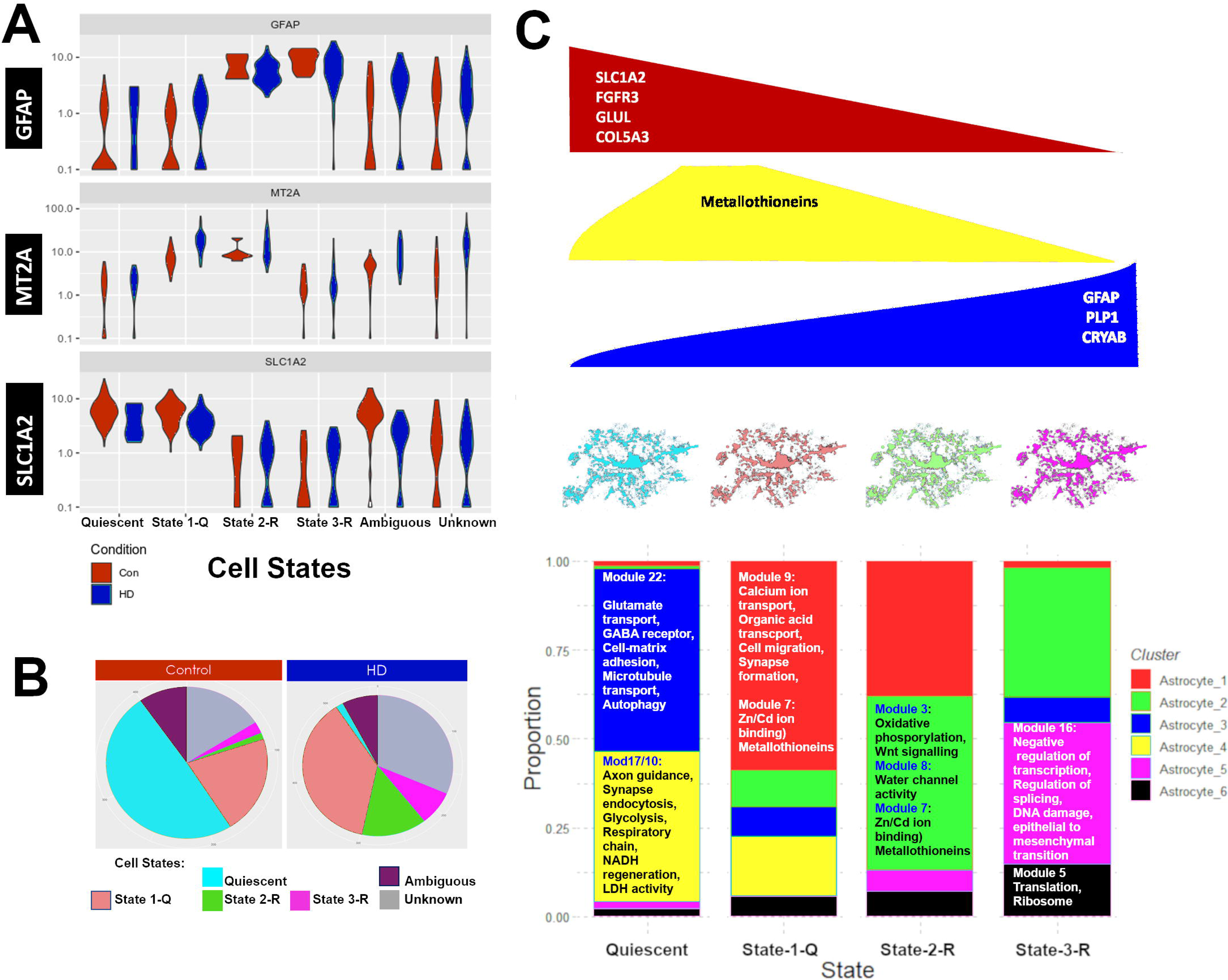
Three reactive astrocytic states in HD. **A**) Supervised classification of astrocytic nuclei based on normalized expression levels of GFAP, MT2A, and SLC1A2 in control (red) and HD (blue) presented as violin plots. Four states are noted: Quiescent, state1Q, state-2R, and state-3R. Ambiguous and unknown state represent cells that met more than one classification condition or none, respectively. **B**) Pie charts of the relative proportions of the astrocytic states in control and HD. **C**) Cartoon summary of the astrocytic states color coded as in (**B**) with respect to the expression of reactive genes such as CRYAB, GFAP, and MTs, as well as protoplasmic astrocyte genes such as SLC1A2, FGFR3, and GLUL. The lower panel shows the proportion of astrocytic clusters colored as described in the legend on the right (same as Figure 3A-C) in each of the quiescent and reactive states (as described in panels A-B). Top gene modules that characterize each astrocytic cluster are described.

A summary of gene expression differences between astrocytic clusters in different states of reactivity summarized as modules is shown in **Figure 6C**. Control astrocytic clusters Astrocyte_3 and Astrocyte_4 are represented in quiescent and state1-Q, and are enriched in genes described by modules 22, 28, 10, and 17, which harbor genes involved in glutamate transport, GABA receptors, autophagy, glycolysis, cell respiration, and synaptic function. HD cluster 1 is represented in states 1-Q and 2-R, while HD cluster 2 is represented in states 2-R and 3-R. Both clusters are enriched in genes of module_7 (*MT*s). Conversely, HD cluster 5is represented mainly in state 3-R and is enriched in module16, harboring genes involved in mesenchymal differentiation, DNA damage, and negative regulation of transcription. Overall, we present complementary views to understand gene expression changes in HD astrocytes in the cingulate cortex, using unsupervised clustering, supervised classification, and gene network analysis.

### Is HD astrocyte pathology cell-autonomous?

Whether the astrocytic reactivity in HD is a cell autonomous reaction or a secondary reaction to neurodegeneration, or both, is yet to be determined. We provide evidence that HTT indeed accumulates in astrocytes in the HD cortex. These HTT aggregates are seen in a minority of HD astrocytes, primarily in layers V and VI of the cingulate cortex, where neurodegeneration is most pronounced [13]. We also provide evidence that neuronal loss and dysfunction is evident in the HD cingulate cortex (See **Supplement**)

Thus, astrocytic reactivity in HD may indeed be secondary to neuronal loss. However, the astrocytic dysfunction may also contribute to neuronal loss and dysfunction. HD transgenic mouse studies that target *mHTT* to astrocytes exhibit neuronal loss, suggesting that mutant HTT expressed in astrocytes alters homeostatic functions and precipitates neuronal dysfunction and loss. Restoration of astrocytic function ameliorates the HD phenotype and prolongs survival *in vivo* [56]. Additional evidence from the HD mouse model R6/2 indicates that chimeric mice engrafted with control human glia including astrocytes exhibited slower disease progression. Conversely, control mice engrafted with HD glia including astrocytes exhibited impaired motor coordination [4]. Thus, it is possible that astrocytic dysfunction in HD maybe partly cell autonomous and partly secondary to neuronal dysfunction. Future studies to correlate the cortical locale of reactive HD astrocytes with the transcriptional phenotype may also help to answer this question.

Our data on neuronal gene expression profiles on the bulk RNAseq level and the single cell level as well as neuronal estimates show major transcriptional and cellular alterations. First, we show that large (pyramidal) neurons are depleted in the HD cingulate (as verified by histopathologic quantification). Second, we show that neuronal genes are downregulated in bulk RNAseq HD samples. For example, pathways involved in glutamate, GABA, neuropeptide Y neurotransmission were downregulated in HD. Finally, our snRNAseq show that glutamatergic neurons, in contrast to interneurons, cluster separately between control and HD. Moreover, neurons upregulate metallothioneins and pathways involved in heat-shock response, voltage-gated ion channels, and protein misfolding. Therefore, neuronal loss and dysfunction is evident in the HD cingulate cortex and may underlie astrocytic reactivity, but it could also be at least partially secondary to astrocytic dysfunction.

### Upregulation of Inflammatory genes in the HD cingulate cortex

The bulk RNAseq analysis showed upregulation of a number of inflammatory genes (see **Supplementary data** and **Supplementary Figure 6**). Our findings highlight the activation of the innate immune system including toll-like receptor pathways. Multiple interleukin signaling pathways were enriched in HD including IL-4, IL-13, and IL-10. Of note, we found IL-10 to be significantly upregulated in HD versus controls. IL-10 has been previously reported to be elevated in the serum and CSF of HD patients [5].

## Conclusions

In summary, we show using snRNAseq from post-mortem human HD cingulate cortex, that astrocytic reactivity can be described in three states with different levels of *GFAP*, metallothionein genes, and quiescent protoplasmic genes. Immune pathways were prominently enriched in the HD cortex, although we were only able to detect them using bulk RNAseq. There were significant gene expression differences between HD and control neurons, particularly in the excitatory neuron populations, along with neuronal loss. We have made our snRNAseq data available using an interactive web application found here (https://vmenon.shinyapps.io/hd_sn_rnaseq/).

## Data availability

Data for bulk RNAseq analysis is provided as a csv file with raw and normalized counts (Supplementary data). Data for snRNAseq can be queried using an interactive web app: https://vmenon.shinyapps.io/hd_sn_rnaseq/. All other data is available upon reasonable request.

## Supporting information

Discription of supplement

## Acknowledgements

This work was funded by the Hereditary Disease Foundation, Taub Institute for the Aging Brain, and the Department of Pathology and Cell Biology at Columbia University. This research was supported by the Digital Computational Pathology Laboratory in the Department of Pathology and Cell Biology at Columbia University Irving Medical Center. We thank Sufian Al-Ahmad and Ahmad Aby-Aysheh from Al-Fardthakh for support with aspects of bulk RNAseq data analysis, Dr. Claudia Deoge for help with RNAscope, and the immunohistochemistry core in the Department of Pathology and Cell Biology at Columbia University Irving Medical Center for help with GFAP-HTT immunostains. We are grateful to the Hereditary Neurological Disease Centre, Wichita, KS, for sending tissues of their HD patients. Conflict of Interest: The authors declare that they have no conflicts of interest.

## Supplementary Data

### Supplementary Methods

#### Quantification of cortical thickness and neuronal densities and statistical analysis

To quantify cortical thickness, the boundary between the grey and white matter was first identified by a neuropathologist (OAD, JEG) on Hematoxylin and Eosin histochemical stain, Cresyl violet histochemical stain, and CD44 immunohistochemical stain. Whole slides were scanned and the images were analyzed in the Aperio™ eSlideManager. A set of lines orthogonal to the pial surface were drawn from the surface to the white matter, and the lengths of these lines were measured. The cingulate cortex was divided into three adjacent regions; abutting the corpus callosum, the convexity of the cingulate gyrus facing the contralateral gyrus, and the sub-frontal surface of the dorsal side of the cingulate gyrus. Three to five measurements were taken per region. Areas with staining artifact or tissue folding were excluded. The average cortical thickness for each region was calculated per case per stain. The number of cases examined was as follows: 4 control and 5 HD for CD44 immunostain, 6-9 HD and 6-8 control for H&E, and 5-8 HD and 6-7 control Cresyl violet. Unpaired t-test with unequal variance was performed to test significance.

For quantification of nuclear densities in whole slide images of Cresyl violet stained sections scanned at 20X, we used a semi-automated method using Qupath v0.2 (Bankhead P et al. 2017). We employed a multi-step approach to first count cells in a ROI, agglomerate (bin) cells counts based on cell area, divide by total count, and finally express these counts as proportions of cells in given area range to the total count. In brief, images were loaded under the “bright field (other)” setting. The staining vector was estimated for each image from a representative region of interest (ROI), excluding unrecognized colors. The Cresyl violet signal was largely identified in the range of Hematoxylin. Next, a script to automate watershed cell detection was used with the following parameters based on “Optical density sum”: Background radius 8□m, Sigma 1.5□m, maximum area 600□m^2^, Threshold 0.2, smooth cell boundaries = True, and separate by shape = False. The results were collected for 3-6 ROI per image. Areas with tissue folds, staining artifact, or blood vessels were excluded. One image from each patient was used. The results were loaded in R v3.6. An empirical threshold of the mean nuclear hematoxylin intensity and sum nuclear hematoxylin intensity was determined to exclude cells/events that represented tangential cuts through cells or background noise. The mean nuclear sum and mean hematoxylin intensity were first normalized. The threshold was determined based on inspecting values from 4 images (and validated in the remaining images). All events were first filtered by the threshold determined for the sum of the hematoxylin intensity, followed by the threshold determined for the mean nuclear hematoxylin intensity. Next, the events of nuclear area (Hematoxylin) were tallied and grouped based on 5 quantiles based on the range of nuclear areas. The counts per case were normalized by the total event count per case to get the relative proportions of nuclei in each of the 5 quantiles of area ranges and analyzed by a two-way ANOVA with an unbalanced design (n= 9 for control and 8 for HD), and ANOVA type set at “III”. The test was performed in R, with the counts per area range as the dependent variable, and the area ranges and Conditions as explanatory variables, with an interaction term between area ranges and Condition. Tuckey test for posthoc testing was done and statistical results were only reported for results of the interaction comparisons.

For quantification of GFAP + cell densities in GFAP/HTT double stained or GFAP single immunostained slides, we counted cells with an area exceeding an empirically-determined threshold, which corresponds to astrocytes with increased GFAP-immunopositive cell area-associated with reactive astrocytosis. The following script was used in QuPath v0.2: runPlugin(’qupath.imagej.detect.cells.WatershedCellDetection’, ‘{“detectionImageBrightfield”: “Optical density sum”, “requestedPixelSizeMicrons”: 0.5, “backgroundRadiusMicrons”: 30.0, “medianRadiusMicrons”: 2.5, “sigmaMicrons”: 2.0, “minAreaMicrons”: 40.0, “maxAreaMicrons”: 600.0, “threshold”: 0.8, “maxBackground”: 3.0, “watershedPostProcess”: false, “excludeDAB”: false, “cellExpansionMicrons”: 7.672413793103447, “includeNuclei”: true, “smoothBoundaries”: true, “makeMeasurements”: true}’). For DAB stained slides, the following empirically determined setting were used: runPlugin(’qupath.imagej.detect.cells.WatershedCellDetection’, ‘{“detectionImageBrightfield”: “Optical density sum”, “requestedPixelSizeMicrons”: 0.5, “backgroundRadiusMicrons”: 10.0, “medianRadiusMicrons”: 1.5, “sigmaMicrons”: 1.5, “minAreaMicrons”: 40.0, “maxAreaMicrons”: 600.0, “threshold”: 0.4, “maxBackground”: 2.0, “watershedPostProcess”: false, “excludeDAB”: false, “cellExpansionMicrons”: 7.67, “includeNuclei”: true, “smoothBoundaries”: true, “makeMeasurements”: true}’). The counts and areas of regions of interests were entered into a Table into R. Cell densities were calculated. Mann-Whitney U test (one-sided) was used to calculate statistical significance.

#### Quantification of GFAP and MT optical density

Quantification of the levels of GFAP and MT immunoreactivity was performed on the images obtained with confocal microscope from triple (GFAP/MT/ALDH1L1) stained sections. Images (merged from the z-stacks of adjacent 6 optical planes [1024 × 1024 pixel resolution, observed area 606 × 606 μm of a 40x field] captured at a z plane distance of 0.4 μm from each other) were transferred into Image J (public domain), grayscaled for each channel, and optical density (OD) was determined in a circle area (diameter 30 μm) centered at the cell nuclei. Only astrocytes with clearly outlined nuclei profiles (DAPI staining) were taken into consideration. At least 5 images from randomly selected gray matter areas from 3 cases of HD and 3 cases of control cingulate cortex were used for quantification. A total of 79 cells (HD) and 49 (control) cells were analyzed. An unpaired two-tailed t-test was used for statistical comparison.

### Supplementary Results

#### Neuronal loss and transcriptional alterations in the HD cingulate cortex

Here we present neuronal data, since neuronal pathology in HD has been the major object of most studies. First, to determine the extent of neurodegeneration in our samples of cingulate cortex, we examined the cortical thickness in different histochemical stains of control and HD tissue. The results showed no significant difference in cortical thickness of the anterior cingulate gyrus between control and HD cases as quantified with a CD44 immunohistochemical stain (**Supplementary Figure 1A-B**), an H&E histochemical stain (**Supplementary Figure 1A, 1C-D**), and a Cresyl violet histochemical stain (**Supplementary Figure 1C, 1E-F**). Next, we quantified the relative proportions of nuclei of different sizes in Cresyl Violet stained sections of the cingulate cortex (**Supplementary Figure 4A**). We categorized the nuclear areas into 5 quantiles. Larger nuclear areas corresponded to pyramidal neurons, whereas smaller areas were either glial or small neuronal nuclei (**Supplementary Figure 4B**). We next conducted a two-way ANOVA analysis of the proportion of nuclei in each area bin between control and HD. Although there were no overall significant effects of the Condition (F ratio: 1.2036 and p value 0.317) or Area range (F ratio 0 and p value 1), there is significant crossover interaction (F ratio of 17.635 and p value 4.72e-10). The proportion of nuclei in an area range (bin) is the opposite, depending on the condition (**Supplementary Figure 4C**). More specifically, the proportion of nuclei within a 13.5-30.8µm^2^ range were increased in HD (Adjusted p value 1.7 e-4), while the proportion of nuclei with areas greater than 104 µm^2^ were decreased in HD compared to control (Adjusted p value 7.6 e-6 – **Supplementary Figure 4C**). The increased proportion of small nuclei could reflect a shrinkage of larger cells or a greater proportion of small, glial cells, or both. We have independent evidence that the proportion of cells of the oligodendrocyte lineage is increased, and is accompanied by a shift to less mature oligodendrocytes (manuscript in preparation).

We first analyzed our bulk RNASeq data, examining the gene ontology terms and Reactome pathways enriched in genes downregulated in HD. The majority of terms relate to neuronal identity (synapse, neuron part, neuron projection, axon part, dendrite tree, and growth cone, for eg.) or neuronal function (GABA receptor complex, regulation of catecholamine secretion, dopamine transport, AMPA glutamate receptor activity, and neuropeptide receptor activity). These GO terms are presented in **Supplementary Figure 4D**. Together, these data suggest that neuronal numbers and function are reduced in these HD cingulate cortex samples.

We then analyzed snRNASeq findings. A total of 1866 neuronal nuclei passed quality control. When neuronal nuclei were placed in a tSNE plot, we distinguished 9 different neuronal clusters (**Supplementary Figure 5A**). HD clusters and control clusters were largely separated, with some overlap (**Supplementary Figure 5B**). A differential gene expression analysis (volcano plot) revealed a number of up- and down-regulated genes in comparing HD neurons with control neurons. The former included MT and heat shock protein genes, as we found in astrocytes, the latter included somatostatin and NPY (**Supplementary Figure 5C**). Analysis of enriched gene ontology terms and Reactome pathways showed the top gene ontology molecular function terms of genes increased in HD included terms related to MTs, metal binding, heat-shock factor 1 (HSF-1) dependent transactivation, misfolded protein binding, molecular chaperones, and, interestingly, voltage gated ion channel activity (**Supplementary Figure 5D**). Gene ontology terms of genes decreased in HD included those related to translation initiation and elongation, selenoamino acid metabolism, regulation of SLITs and ROBOs, structural constituents of ribosomes, and neuropeptide receptor binding (**Supplementary Figure 5E**). A list of differentially expressed genes in HD versus control neurons as well as the full results of GO term enrichment analysis are provided in **Supplementary Table 6B-C**. A list of the top neuronal cluster markers is provided in **Supplementary Table 7**.

Differential gene expression of the neuronal clusters was displayed in a heat map (**Supplemental Figure 5F**). Clusters 1, 2, and 3 show high expression of genes characteristic of inhibitory interneurons (for example, *VIP*, *GAD1*, *CALB2*, *SST*). Clusters 4-9 are relatively much lower in these transcripts, and higher in a set of other transcripts (for eg. *SLC17A7*, *NGRN*, *CHN1*, *SNCA*, *NEFL*, *GRIN1*, *TMSB10*, *BASP1*), associated with excitatory neurons. However, an inspection of the heat map does show gene expression differences that vary among these clusters, suggesting significant variation in populations of excitatory neurons. The clusters that correspond to inhibitory neurons contain mixtures of cells from control and HD brains, while clusters from excitatory neurons are divided up much more clearly between control and HD brains. This suggests that changes in excitatory neuron gene expression is more affected by HD than inhibitory neuron gene expression. We validated that a number of the transcripts that we found associated with the neuronal clusters were indeed expressed by neurons by referring to *in situ* hybridizations from the Allen Brain Atlas (**Supplementary Figure 5G**).

#### A subset of Microglial genes is captured by single nuclei RNAseq in the cingulate HD cortex

Because of the inflammatory gene expression in HD and because both astrocytes and microglia contribute to inflammation, we are presenting findings on microglial gene expression. Investigation of microglial gene expression in bulk RNAseq data based on a list of specific microglial genes from Patir et al [40] showed a significant increase in multiple immune genes (**Supplementary Figure 6A**). Multiple immune pathways were enriched in HD including IL-4, IL-10, and IL-13 signaling (**Figure 1D**). There is significant heterogeneity between HD cases; with the signature being largely driven by two HD cases (5300 and 5575). The genes increased in HD included *TLR*, *HLA*, Fc Gamma receptors, and complement factor genes. We investigated the individual transcriptional profiles from snRNAseq of HD cases 5575 and 5493 and controls 5382 and 5404. We identified 147 nuclei as microglia (3.7% of total nuclei, **Supplementary Figure S6B**). Control and HD nuclei clustered separately (**Supplementary Figure S6C**). Differential gene expression analysis showed 148 genes to be differentially expressed at a false discovery rate of 0.25 (**Supplementary Figure S6D-E**). From the differentially expressed genes, seven genes were found to be also upregulated in the bulk RNAseq data (*SPP1*, *CD163*, *SLC11A1*, *VSIG4*, *HLA-DRB5*, *FCGR3A*, *MS4A6A*). Three genes were downregulated compared with the bulk RNAseq (*CSF3R*, *CSF2RA*, and *TREM2*). Note this data reflect gene expression in nuclei from two HD cases and two controls, as opposed to n=6 per group in bulk RNAseq. The results indicate that with our approach, detecting microglial gene alterations at the single nucleus level might not be as sensitive as bulk RNA sequencing. The genes used in the heatmap of **Supplementary Figure 6** are provided in **Supplementary Table 7**.

A limitation we note in our study is that we detected only a small number of microglial nuclei. The paucity of microglial nuclei and the sparsity of nuclear reads likely explains why we did not detect many differentially expressed microglial genes. In addition, only one of the two HD donor brain samples showed large upregulation of microglia reads in the bulk RNAseq analysis. Thus, our approach may not be powered to illuminate transcriptional alterations of microglia at the single nucleus level. That being said, we still found alterations of the immune system using the bulk RNAseq data.

#### HD modifier genes expression in neurons and astrocytes

We investigated the expression of HD modifier genes in neurons and astrocytes (Gem-HD consortium, 2015). We note that while there were no detected changes in the expression of HTT between control and HD neurons, HD astrocytes overall expressed lower levels of HTT (log fold change = −0.439, FDR=0.0189). Next, we turned our attention to neurons. We detected no significant changes in the expression of HD modifier genes *FAN1*, *RRM2B*, or *UBR5* in HD neurons versus control (**Supplementary table 6A**). Conversely, the expression of *MLH1* was significantly reduced in HD neurons compared with controls (log fold change = −0.449, FDR = 2.99 E-5; **Supplementary Table 6A**). The expression of *MLH1* was mostly noted in the control neuronal clusters 7 and 8 (excitatory neurons – data not shown). The picture is different in astrocytes (**Supplementary Tables 2C).** We found, the expression of both *MTMR10* and *UBR5* were reduced in HD astrocytes compared to controls (*MTMR10* log fold change = −0.409, FDR = 0.014, UBR5 log fold change −0.673, FDR = 3.18 E-5). The expression of *FAN1* or *RRM2B* showed no significant difference between HD and control astrocytes. Interestingly, *HTT*, *UBR5*, and *MLH1* were not differentially expressed between HD and controls at our threshold in bulk RNAseq (absolute log fold change levels > 0.5). In contrast, MTMR10 was significantly increased (log fold change = 0.89 and FDR=0.015 - **Supplementary Table 5**). In conclusion, the data reveals significant difference between HD astrocytes and neurons at the single nucleus level; HD astrocytes expressed lower levels of *HTT* and the disease modifiers *UBR5* and *MTMR10*. Conversely, HD neurons did not show significant changes in *HTT*, but showed overall lower expression of the disease modifier *MLH1*. These changes were not reflected in differential gene expression changes in bulk RNAseq, which is a representation of both gene expression per cell type as well as cell type proportions. This may underlie the differences we see in *MTMR10, MLH1, UBR5*, and *HTT* in snRNAseq versus bulk RNAseq.

#### Cell type-specific markers derived from nuclear transcripts

We provided a list of astrocyte sub-cluster markers (“**Consensus clustering using SC3 for sub-clustering”, above**, and **Table 2**). We also derived sets of cell type cluster markers (**Supplementary Table 1C**). For this, we used the cell-type gene lists from neurons, astrocytes, microglia, oligodendrocytes, OPCs, and endothelial cells. Note that there are many overlaps between this list and the original, literature-based, list we used for the Cell Classifier (**Supplementary Table 1B**), but some genes appear in one but not the other. These differences may result from different levels of transcripts in the nucleus vs. cytoplasm and nucleus combined (for single cell RNASeq).

### Supplementary Figure Legends

**Supplementary Figure 1:**
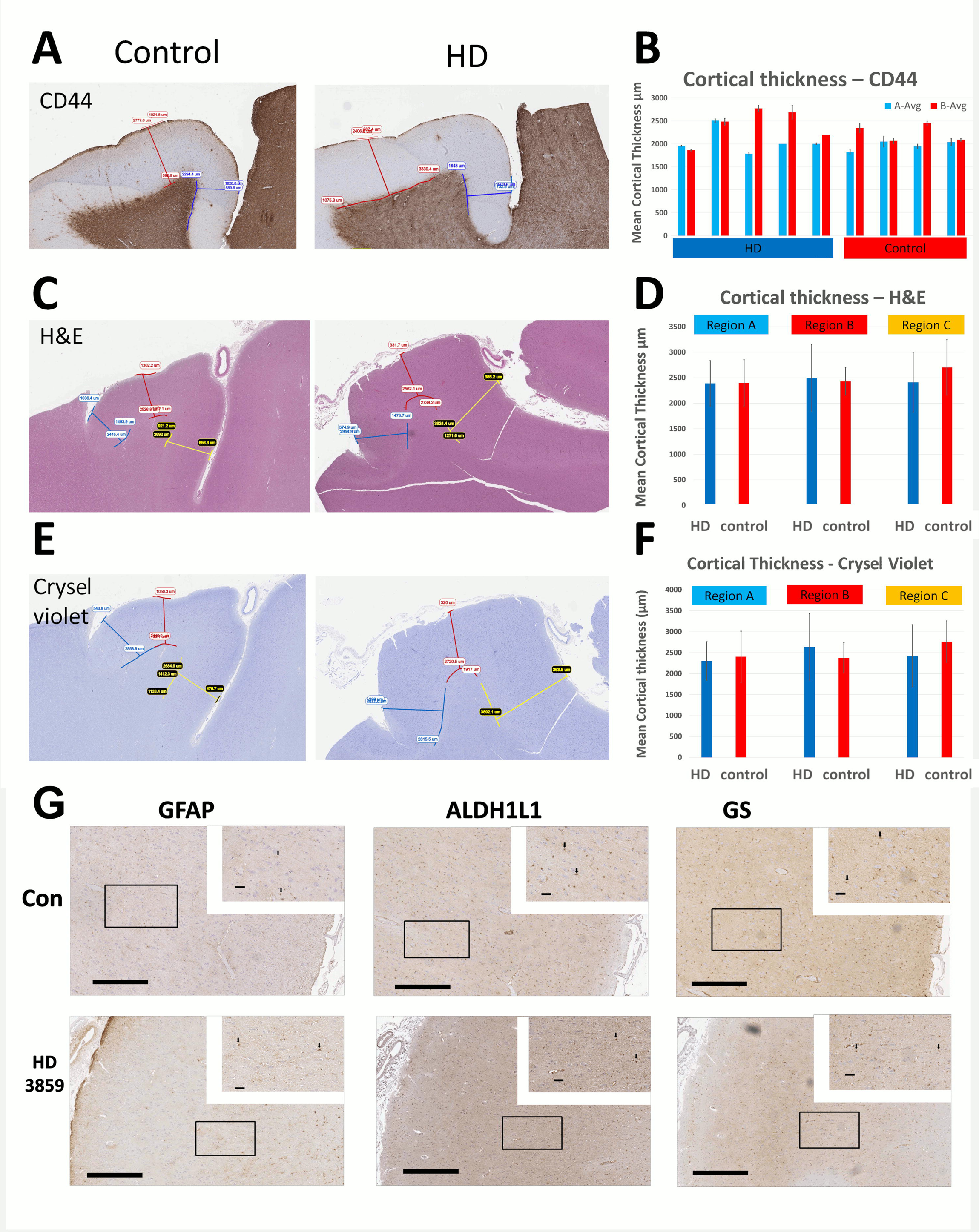
Cortical thickness of the cingulate in HD. Representative images of cortical thickness measurements performed in sections stained for CD44 (**A**), Hematoxylin and Eosin (H&E) (**C**), and Cresyl violet (**E**). Bar graphs showing average cortical thickness in individual cases in in the CD44 immunostain (**B**), with two regions quantified highlighted in blue and red. Bar graphs showing average cortical thickness of control and HD sections stained for H&E (**D**) and Cresyl violet (**F**). The regions quantified in the cingulate cortex are color-coded in the images, which is reflected in the bar graphs. No significant differences were identified between control and HD using unpaired t-tests. N =4 control and 5 HD for CD44 immunostain, 6-9 HD and 6-8 control for H&E, and 5-8 HD and 6-7 control Cresyl violet. **G**) Immunohistochemical staining for GFAP, Glutamine Synthetase (GS), and ALDH1L1 of a representative control and the Juvenile Huntington (T3859). Images are shown at 5X, and insets at μm.

**Supplementary Figure 1: Related to Figure 2.**
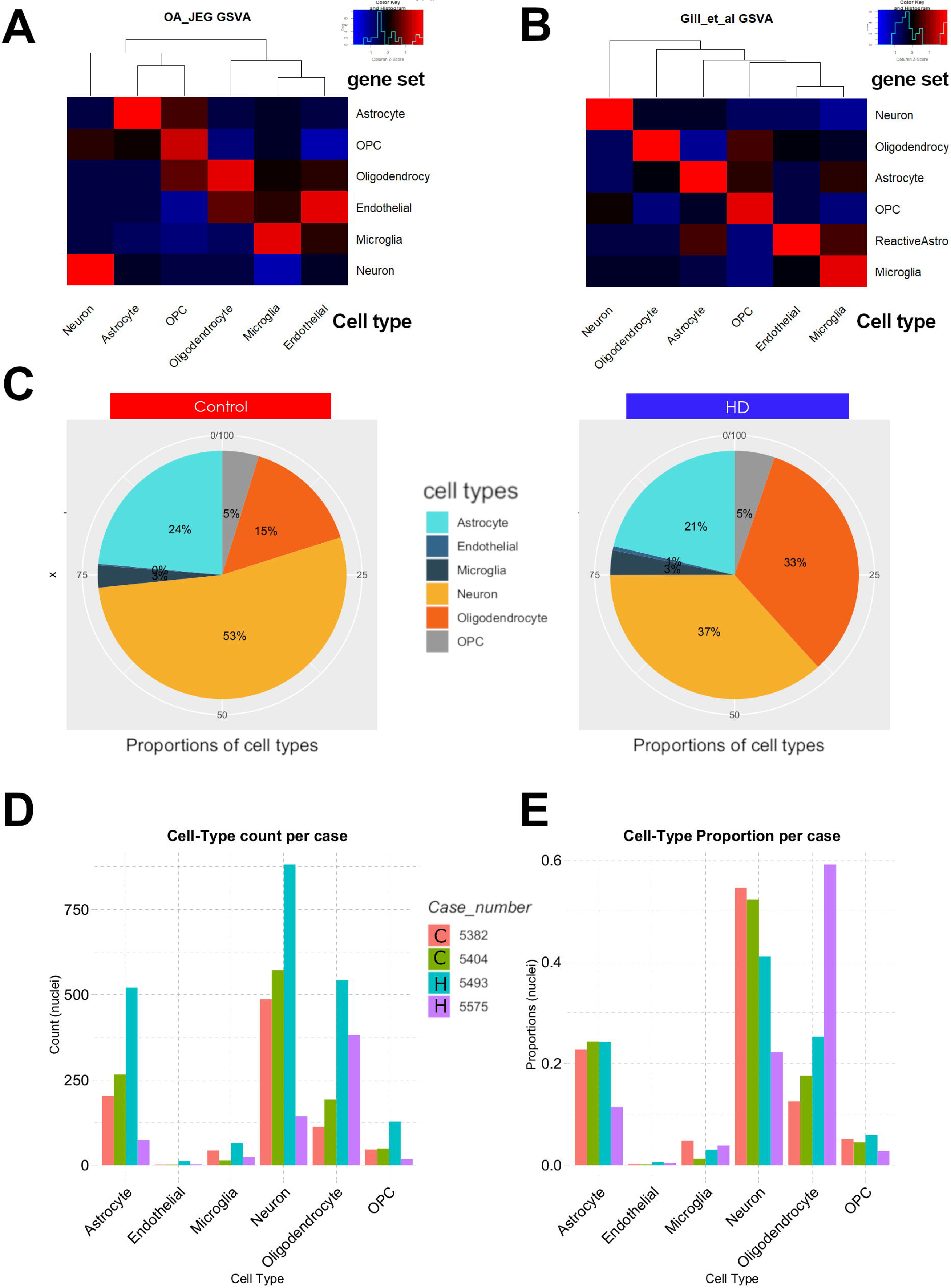
Gene set variation analysis (GSVA) of the average normalized expression of all nuclei in one cell-class/type. Cell-type specific gene sets derived from the literature (**A** OA and JEG) and Gill et al. [10] (**B**) are shown in the rows. Cell-types are shown in columns. The z-scaled enrichment scores of the cell-type averages are shown in the heat maps (**A-B**). The proportions of cell-types in Control (Right) and HD (Left) nuclei. Percentages per cell-type are shown in the pie chart (**C**). Bar-plots of count of nuclei per cell-type per case (**D**). Barplots of the proportions of cell-type per case (C=Control, H=HD) (**E**).

**Supplementary Figure 3:**
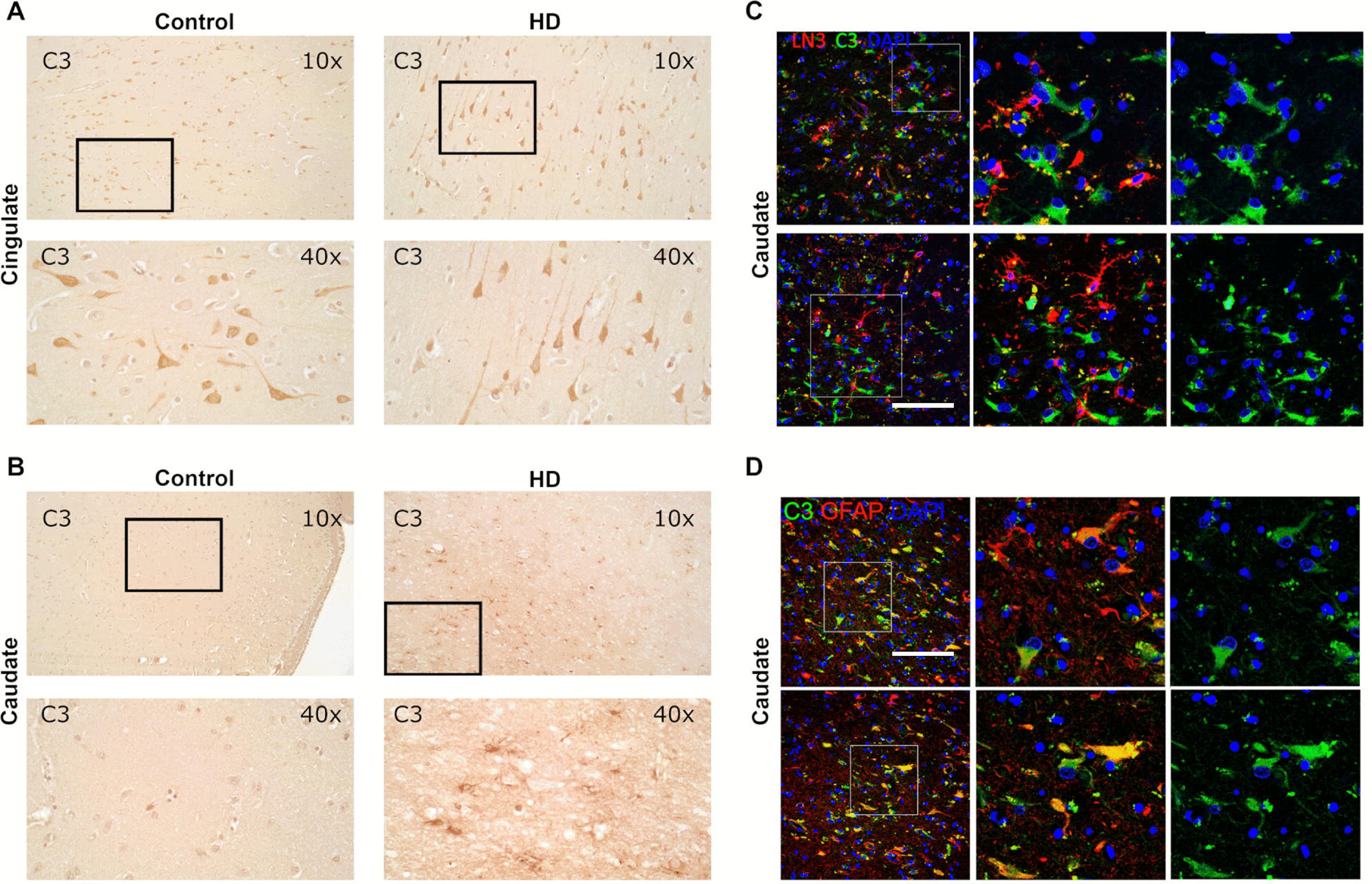
Complement factor 3 (C3) immunostaining in the HD caudate and cingulate. **A-B**) Micrographs of immunostaining for C3 in the cingulate cortex (**A**) and caudate nucleus (**B**) of control and HD grade III/IV taken at 10X (100X total magnification). The boxed areas are shown at 40X in the lower panels (400X total magnification). **C-D**) Dual immunostaining for C3 (green) and GFAP (red -**C**) or LN3 (red – **D**) in the caudate nucleus of a representative HD case (**C**). Nuclei stained with DAPI are shown in blue. Scale bars indicate ####. A total of 3-4 cases per group were examined.

**Supplementary Figure 4:**
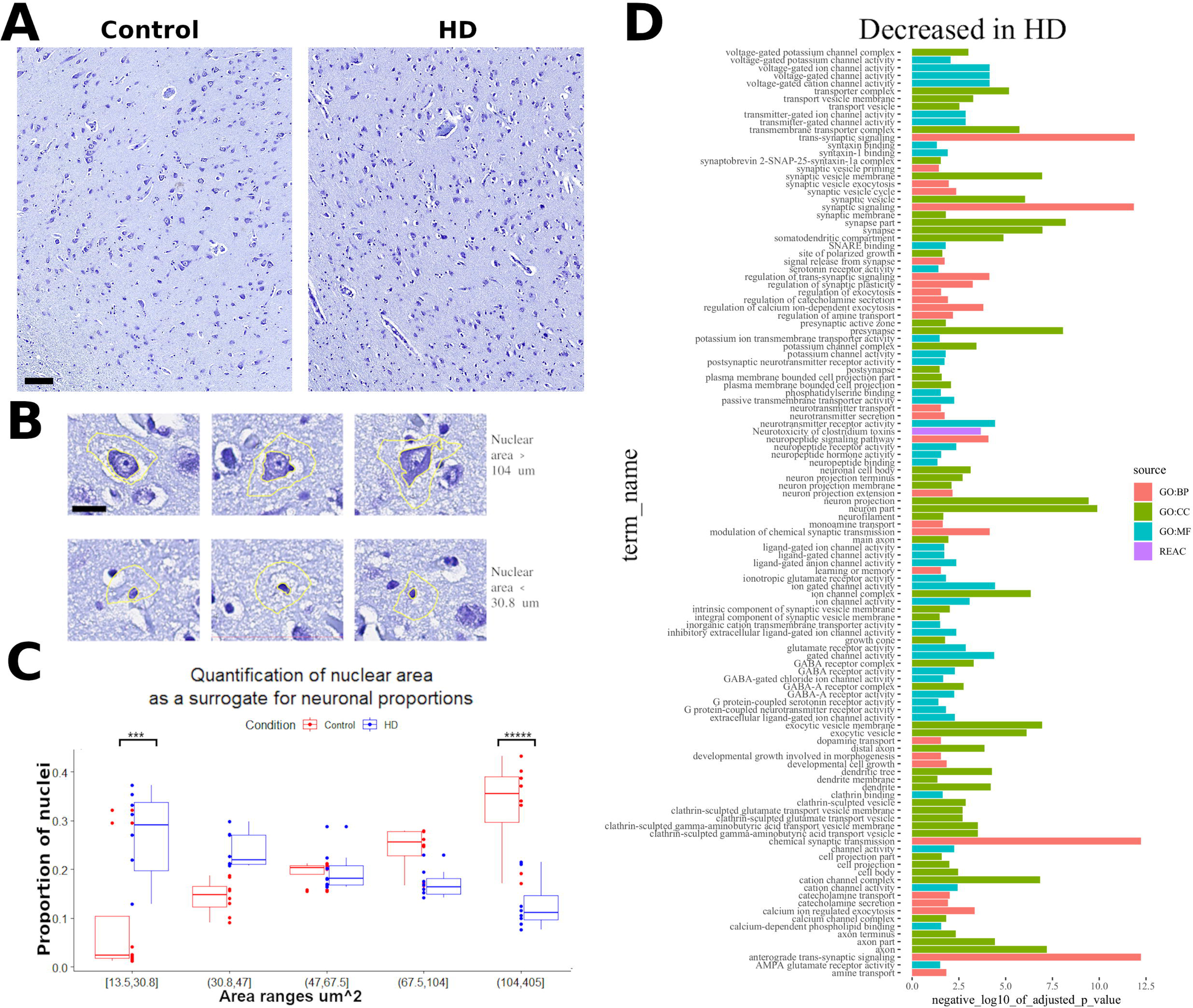
Neuronal loss and dysfunction in the HD cingulate cortex. **A**) Representative images of crystal violet stained cingulate cortex sections from a control case and a grade 4 HD. The subcortical white matter is shown in the lower right corner. Note the relative abundance of the small “glial” nuclei in the HD cortex. Scale bar = 100um. **B**) Representative images of cells with large nuclear area (5^th^ quantile >104 um^2^) in the upper row and cells with small nuclear area (1^st^ quantile <30.8 um^2^) in the lower row. These areas correspond to neurons and glia, respectively. **C**) Quantification of the relative proportions of nuclei quantified (y-axis) within different area ranges (x-axis). Boxplots are shown in addition to points representing individual cases. Clue indicates HD and red controls. N=9 for control N=8 for HD. ***: p value < 0.001, *****: p value < 0.000001. **D**) GO term and Reactome pathway enrichment analysis of genes significantly downregulated in HD in the bulk RNAseq analysis of the cingulate cortex. All of the pathways shown are significantly enriched after Benjamini-Hochberg False discovery rate correction (<0.05). The source of the GO term is color coded. P value of enrichment is represented by the length of the bar per gene ontology.

**Supplementary Figure 5:**
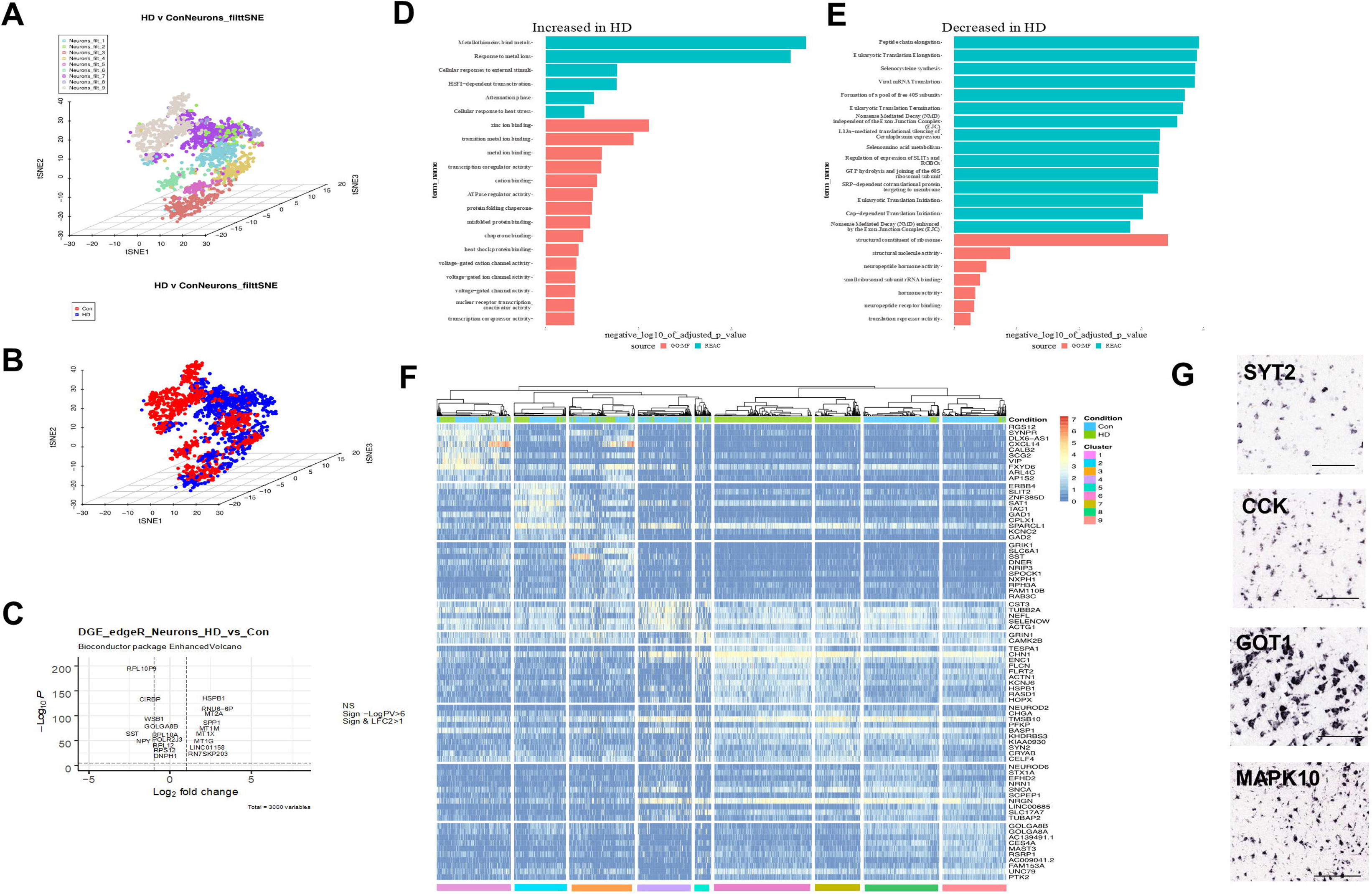
Differential gene expression patterns in neurons. **A)** tSNE plot showing 9 different neuronal clusters. **B)** Here we divided up nuclei in the tSNE plot into HD (blue) and control (red). Some of the clusters appear relatively homogeneous with respect to condition, while others appear more mixed. **C)** Differential gene expression as a volcano plot, showing some of the highly differentially expressed genes. **D)** GO terms and Reactome pathway enrichment analysis of genes significantly increased in HD over all neurons. **E)** GO terms and Reactome pathway enrichment analysis of genes significantly decreased in HD over all neurons. The source of the GO term is color coded. P value of enrichment is represented by the length of the bar. **F)** Gene expression heat map of cluster markers showing nuclei (Columns) and specific genes (Rows). Condition (Con versus HD) and neuronal clusters are color-coded on the top and bottom, respectively. Cluster-specific gene markers were identified using Wilcoxon signed rank test comparing gene ranks in the cluster with the highest mean expression against all others. p-values were adjusted using the “Holm” method. **G)** Examples of *in situ* hybridization of 4 of the neuronal genes (© 2010 Allen Institute for Brain Science. Allen Human Brain Atlas. Available from: human.brain-map.org). Scale bars: GOT1 100□m, others 200□m.

**Supplementary Figure 6:**
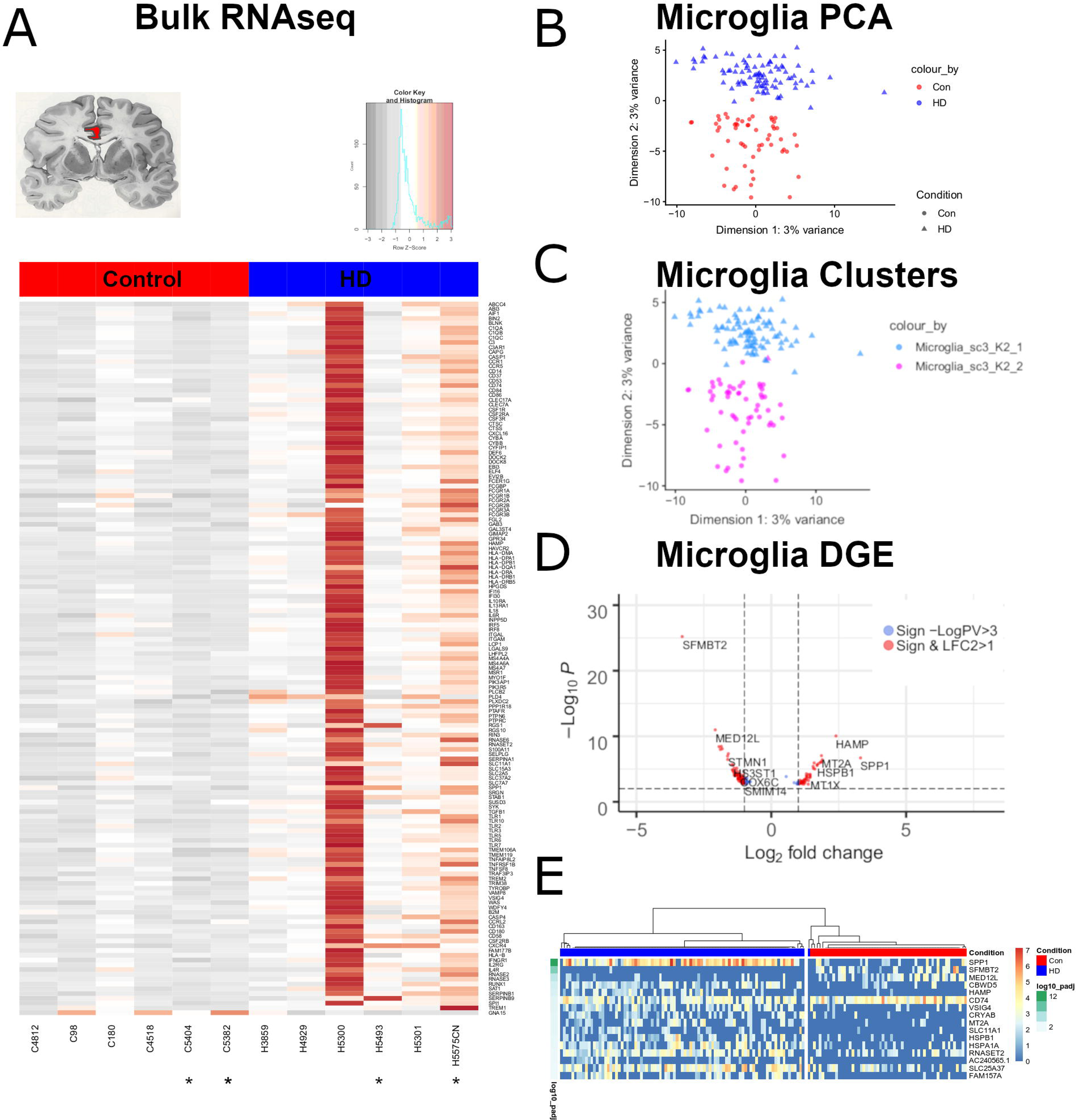
Microglial gene expression alterations in the cingulate cortex. **A**) Differential gene expression heatmap showing a subset of significantly differentially expressed microglial genes in control and HD cingulate cortex – cingulate cortex shown in red in the top right pictogram (Microglial gene list was adapted from Patir et al [40]). **B**) Principle component analysis plot of microglia nuclei. Control nuclei are shown in red, HD in blue. **C**) Clustering of control (circles) and HD (triangles) nuclei shows they cluster separately. Colors denote clusters as determined by SC3 consensus clustering (K=2). **D**) Differential genes expression of between control and HD microglial nuclei displayed as a volcano plot with significance set at p<0.05 – genes with –log10 p value (LogPV) of >3 are shown in blue, and those with -logPV of >3 and log2 fold change >2 are shown in red. Analysis was performed in EdgeR using the likelihood ratio test. **E**) Differential genes expression heatmap between control (red bar) and HD (blue bar) nuclei showing significantly differentially expressed genes (-log10 p value scale shown on the left). Significance was determined using Kruskal-Wallis test (p<0.05).

**Supplementary Figure 7:**
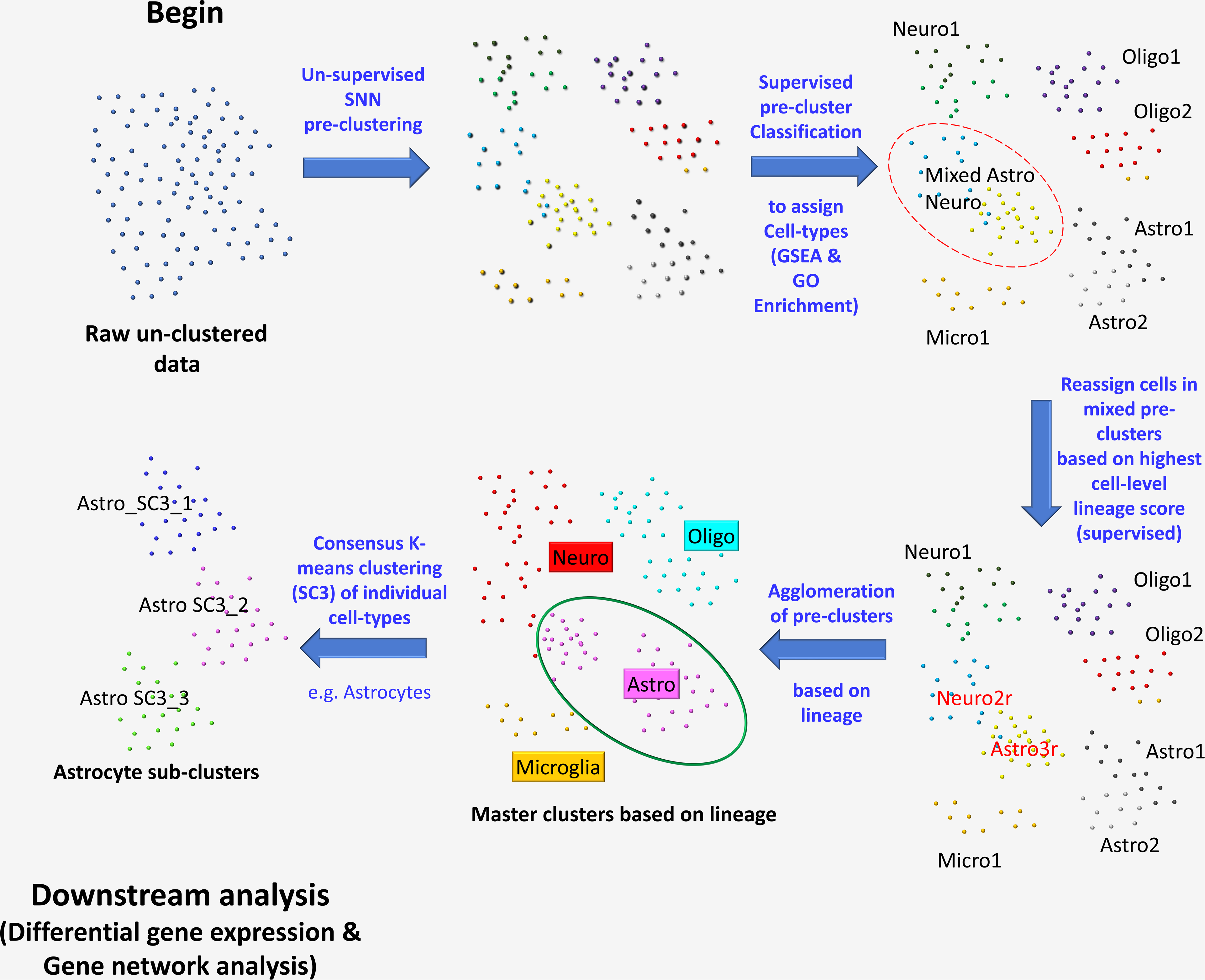
Outline of the snRNAseq analysis pipeline. Briefly, filtered raw un-clustered data is pre-clustered using a shared nearest neighbor algorithm. The pre-clusters are classified into cell classes/lineages using gene set enrichment analysis for specific lineage genes and examining GO terms of the top pre-clusters markers. Next, mixed pre-clusters are identified and the cells in these pre-clustered are re-classified based on the cell-specific lineage-scores (Cell classifier tool). Next, clusters of the same lineage are agglomerated and cell-classes/lineages are analyzed in isolation from the remaining cell classes using SC3 consensus clustering into sub-clusters (Astrocyte sub-clusters are shown as an example). These sub-clusters are used for downstream analysis.

**Supplementary Figure 8:** Validation of astrocytic sub-clusters. Confocal Immunofluorescence images of the cingulate cortex of an HD brain. GFAP (green), ALDH1L1 (red), MT (cyan), DAPI (white) are shown. A merged panel is displayed on the right. The arrow indicates a GFAP+ALDH1L1+MT+ astrocyte (example of Astrocytic Cluster 2). The arrowhead indicates a GFAP-ALDH1L1+MT+ astrocyte (example of Astrocytic Cluster 1). Scale bar = 20□m. Single optical planes are shown.

## Notes

https://vmenon.shinyapps.io/hd_sn_rnaseq/

