## Supplementary material for "Single-nucleus RNA-seq identifies Huntington disease astrocyte states": Discription of supplement

Supplementary movie 1: Three dimensional view of figure 3b
Supplementary table 1: Genes used in gene set enrichment analysis (Gill et al 2014), in cell classifier, and discovered cell-type marker genes based on single nuclei RNAseq data

Supplementary table 2: Differential gene expression between control and HD Astrocytic nuclei, GO term enriched in top differentially upregulated and downregualted genes

Supplementary table 3: Differential correlation analysis between astrocytic genes in HD and control

Supplementary table 4: Astrocytic gene co-expression modules, module information, module genes, GO term enrichment in modules

Supplementary table 5: Results of differential gene expression analysis of Bulk RNAseq controled for age and gender with log fold change threshold of 1.5

Supplementary table 6: Differential gene expression between control and HD neuronal nuclei, GO term enriched in top differentially upregulated and downregualted genes

Supplementary table 7: Microglil genes shown in heatmap of figure S6A

Supplementary data: 
1: Raw counts and RPKM counts of Bulk RNAseq data
2: Results of differential gene expression analysis of each astrocytic cluster against all other clusters  
